# Spatiotemporal heterogeneity of glioblastoma is dictated by microenvironmental interference

**DOI:** 10.1101/2021.02.16.431475

**Authors:** Vidhya M. Ravi, Paulina Will, Jan Kueckelhaus, Na Sun, Kevin Joseph, Henrike Salié, Jasmin von Ehr, Lea Vollmer, Jasim K. Benotmane, Nicolas Neidert, Marie Follo, Florian Scherer, Jonathan M Goeldner, Simon P. Behringer, Pamela Franco, Ulrich G. Hofmann, Christian Fung, Jürgen Beck, Roman Sankowski, Marco Prinz, Saskia Killmer, Bertram Bengsch, Axel Karl Walch, Daniel Delev, Oliver Schnell, Dieter Henrik Heiland

**Affiliations:** Microenvironment and Immunology Research Laboratory, Medical Center, University of Freiburg, Germany; Department of Neurosurgery, Medical Center, University of Freiburg, Germany; Faculty of Medicine, Freiburg University, Germany; Neuroelectronic Systems, Medical Center, University of Freiburg, Germany; Translational NeuroOncology Research Group, Medical Center, University of Freiburg, Germany; Neurosurgical Artificial Intelligence Laboratory Aachen (NAILA), Department of Neurosurgery, RWTH University o f Aachen, Aachen, Germany; Research Unit Analytical Pathology, Helmholtz Zentrum München, Neuherberg, Germany; Department of Medicine II: Gastroenterology, Hepatology, Endocrinology, and Infectious Disease, University Medical Center Freiburg, Freiburg, Germany; Department of Medicine I, Medical Center – University of Freiburg, Faculty of Medicine; Department of Neurosurgery, RWTH University of Aachen, Aachen, Germany; Center for NeuroModulation (NeuroModul), University of Freiburg, Freiburg, Germany; Institute of Neuropathology, Medical Center - University of Freiburg; Signalling Research Centres BIOSS and CIBSS, University of Freiburg, Germany

## Abstract

Glioblastomas are highly malignant tumors of the central nervous system. Evidence suggests that these tumors display large intra- and inter-patient heterogeneity hallmarked by subclonal diversity and dynamic adaptation amid developmental hierarchies^1–3^. However, the source for dynamic reorganization of cellular states within their spatial context remains elusive. Here, we in-depth characterized glioblastomas by spatially resolved transcriptomics, metabolomics and proteomics. By deciphering exclusive and shared transcriptional programs across patients, we inferred that glioblastomas develop along defined neural lineages and adapt to inflammatory or metabolic stimuli reminiscent of reactive transformation in mature astrocytes. Metabolic profiling and imaging mass cytometry supported the assumption that tumor heterogeneity is dictated by microenvironmental alterations. Analysis of copy number variation (CNV) revealed a spatially cohesive organization of subclones associated with reactive transcriptional programs, confirming that environmental stress gives rise to selection pressure. Deconvolution of age-dependent transcriptional programs in malignant and non-malignant specimens identified the aging environment as the major driver of inflammatory transformation in GBM, suggesting that tumor cells adopt transcriptional programs similar to inflammatory transformation in astrocytes. Glioblastoma stem cells implanted into human neocortical slices of varying age levels, independently confirmed that the ageing environment dynamically shapes the intratumoral heterogeneity towards reactive transcriptional programs. Our findings provide insights into the spatial architecture of glioblastoma, suggesting that both locally inherent tumor as well as global alterations of the tumor microenvironment shape its transcriptional heterogeneity. Global age-related inflammation in the human brain is driving distinct transcriptional transformation in glioblastomas, which requires an adjustment of the currently prevailing glioma models.

## Article

In recent years, novel technologies for single-cell analysis have provided insights into transcriptional regulation and the dynamic evolution of single cells within brain tumors as well as healthy human brain^1,4–9^. Large single-cell RNA sequencing (scRNA-seq) studies of high- and lower-grade glioma have elegantly demonstrated that intratumoral heterogeneity and dynamic plasticity across cellular states are hallmarks of malignant brain tumors^1,7,9^. It was assumed that this dynamic adaptation falls within four different states, namely the mesenchymal-like (MES-like), neural progenitor cell-like (NPC-like), astrocyte-like (AC-like) and the oligodendrocytic precursor cell-like (OPC-like) state, mirroring developmental stages of the human brain^1,3^. Within this complex network of glioma, it was shown that neighboring cells such as neurons, glial- and immune cells contribute to the intricate and dynamically heterogeneous system^1,7,10–13^. However, a major drawback of single cell analysis is the lack of information regarding their spatial arrangement, which allows only indirect predictions of cellular and microenvironmental interactions. The spatial organization of tissue is of high importance in a number of organs, and the brain above all, is particularly dependent on the spatial organization of cortical layers. Thus, it is likely that spatial organization patterns are also imitated by CNS-derived malignancies. Spatial transcriptomics, a novel technology is able to provide transcriptomics data at nearly single-cell resolution, while preserving the spatial architecture^14–16^.

### Deciphering spatially resolved transcriptional heterogeneity and lineages

To characterize the spatial architecture of glioblastoma, we created an atlas of spatially resolved transcriptomics (stRNA-seq) of twenty-four specimens resulting in 94.482 transcriptomes across different age-groups, anatomic regions and pathologies, ***Figure 1a and Extended Data Figure 1-3, Patient information in Supplementary Table 1***. Transcriptomes from non-malignant samples demonstrated similarities across patients whereas malignant transcriptomes were marked by individual gene expression patterns, ***Figure 1b***. To evaluate whether malignant transcriptomes resulted from somatic alterations, we estimated copy number variations (CNVs) from the average expression of genes in large chromosomal regions within each spot, which confirmed the typical gain in chromosome 7 and/or loss in chromosome 10 in the majority of malignant spots, ***Figure 1c-d and Extended Data Figure 1-3***. The high number of individual copy-number alterations and mutational profiles are assumed to drive patient-specific transcriptional regulation^1^ resulting in individual clusters of transcriptomes, similar to results seen in other studies^1,2^.

**Figure 1:**
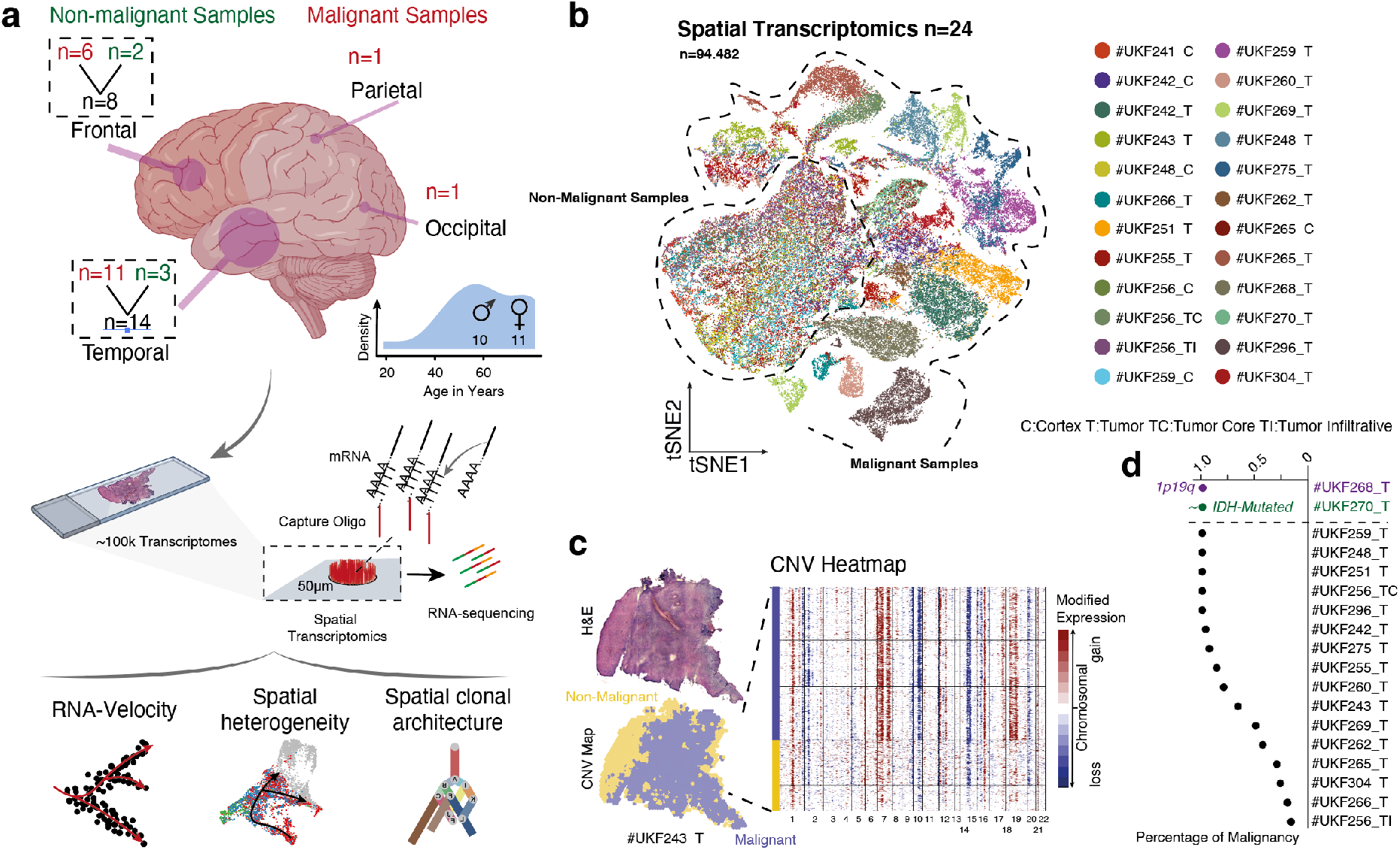
a) Illustration of the workflow and the cohort of spatial datasets (right) and an overview of analytic approaches (right). b) A scatterplot of the tSNE representation with distinct areas of malignant and non-malignant samples. Color reflects individual specimens and patients. c) Illustration of an example of patient #UKF243, a tumor sample which also contained non-malignant areas (marked in yellow) as indicated by the CNV heatmap at the right side. d) Dotplot of the percentage of malignant spots within the stRNA-seq data set based on CNV estimation.

In order to address the intra-tumor heterogeneity with respect to their spatial architecture, we estimated shared signatures across patients which reflect common states and lineages within glioblastoma (GBM), using a combination of two mutually reinforcing approaches. First, we mapped distinct transcriptional programs of individual tumors and then sought for shared programs across all patients. Next, we determined the spatial expression patterns through a generalized linear spatial model and identified recurring patterns across all patients. Through integration of both approaches, we were able to map common transcriptional programs within the spatial context of glioblastoma. After eliminating small and partially overlapping clusters within each patient, we identified a total of 139 patient-specific clusters, ***Extended Data Figure 4a***. To identify shared expression modules, we excluded cell cycle-associated clusters and identified 6 distinct modules that were consistently expressed across all patients, ***Figure 2a*, Supplementary Table 2 and *Extended Data Figure 4a-c***. Of note, this approach allowed us to understand the biological significance of transcriptional programs among patient-specific clusters that occur repetitively and were robustly expressed. Our identified modules encompass two major groups involving developmental and inflammatory/hypoxia-associated transcriptional programs, later referred to as “reactive states”. In contrast to the recent described injury response signature^2^ of glioblastoma, our data indicate that two distinct subtypes of reactive states coexist and emerge spatially segregated from each other, ***Extended Data Figure 5a-b***. The first module revealed a strong enrichment in glycolysis-related pathways and those involved in the response to reduced oxygen-levels (false discovery rate [FDR] < 0.01, hypergeometric test), therefore named as “Reactive Hypoxia” ***Extended Data Figure 5a***. The second module was marked by an enrichment in INF-gamma signaling (false discovery rate [FDR] < 0.01, hypergeometric test), the expression of numerous immune-related genes (e.g. *HLA-DRA, HLA-A, HLA-B*) and the signature genes of inflammatory (also referred to as A1-state^17^) reactive astrocytes (e.g. *GFAP, VIM, CD44*), and is henceforth named as “Reactive Immune”, ***Extended Data Figure 5b***. Spatially resolved projection of both signatures revealed a partial overlap, explained by a subset of genes which were upregulated in response to both reactive signatures such as *CCL2, CHI3L1* and complement factors, ***Extended Data Figure 5c-d***. The remaining modules (3-6) were referred to as “lineage states”, containing genes which were associated with developmental stages, ***Figure 2a-b***. To align our modules along known development hierarchies, we estimated the similarity to gene signatures of developmental cell types^18^, ***Extended Data Figure 5e***, and therefore named the modules as “OPC-like”, “Radial-Glia-like”, “NPC-like-Early Development” and “NPC-like-Late Development”, ***Figure 2b, Extended Data Figure 5f-k***.

**Figure 2:**
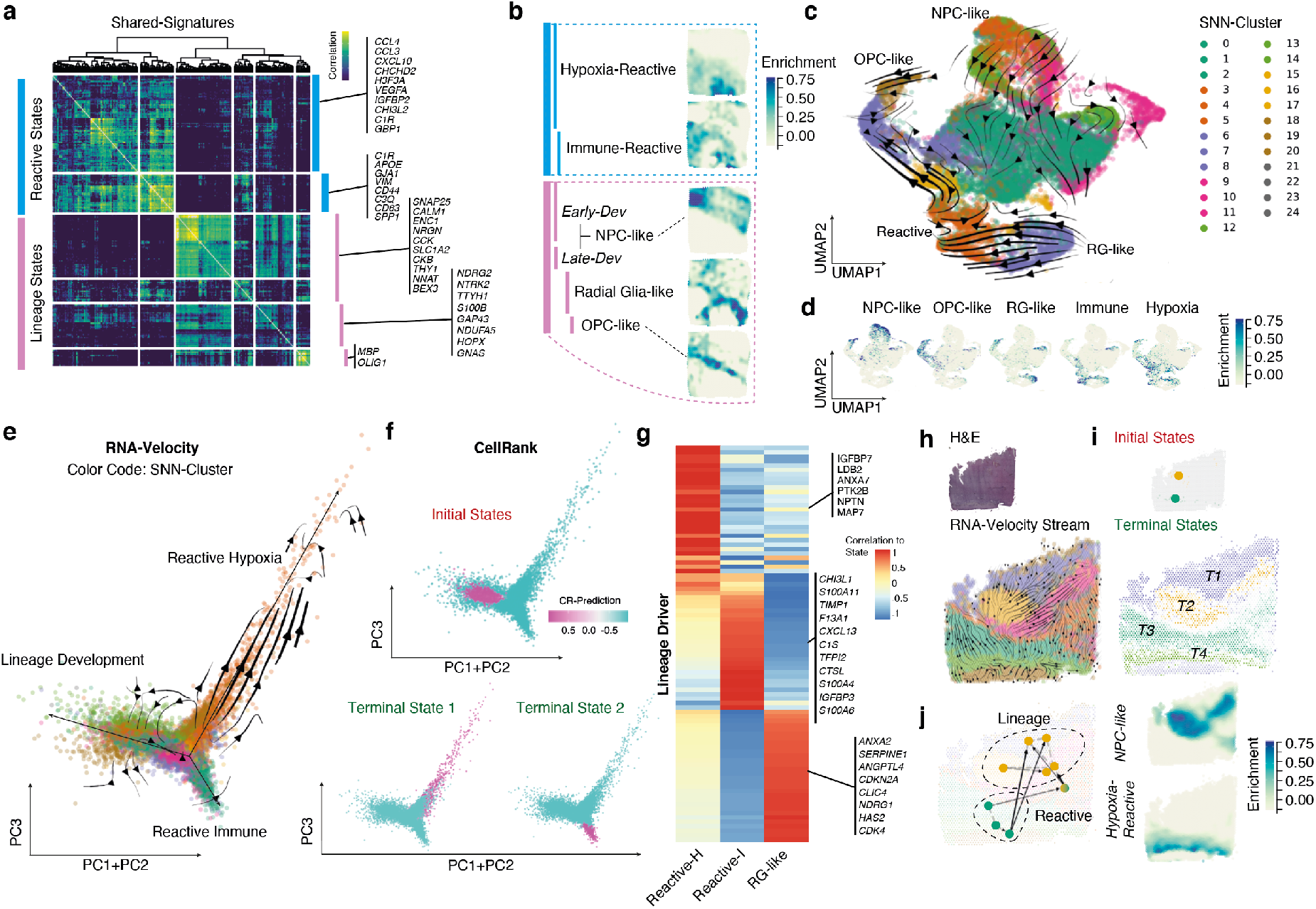
a-b) Correlation heatmap of meta-modules of 6 shared signatures across all tumor spots. Signature genes are listed on the right. The two major subgroups, lineage and reactive states and associated subgroups are illustrated for the #UKF275_T sample (b). c) Dimensional reduction scatter plot (UMAP) and velocity streams called from scVelo are illustrated. Colors indicate the determined cluster (SNN). d) Dimensional reduction scatter plot (UMAP) indicates the enrichment (z-scored) of established signatures (a). e) Scatterplot with the first two eigenvectors on the x-axis and the third eigenvector on the y-axis. Arrows indicate the RNA-velocity streams. f) Estimation of initial and terminal states using CellRank, in a representation similar to (e). g) CellRank based estimation of lineage driver genes are illustrated in a heatmap. h) RNA-velocity streams are presented in space, CNV estimation confirmed the chromosomal alteration of all spots. i) Illustration of the two estimated initial states (CellRank) are demonstrated in the upper panel. RNA-velocity fate mapping determined four terminal states which are presented at the bottom plot. Color density indicates the prediction values for each terminal state. j) Aggregation of individual fate maps into a cluster-level fate map, using partition-based graph abstraction (PAGA) with directed edges, indicates the direction of differentiation at spatial resolution (right panel). Spatial surface plot for the gene set enrichment of NPC-like and reactive hypoxia states (right panel).

Next, we focused on repeating spatial patterns, where we identified a total number of 81 genes, shared across all patients, ***Extended Figure 4e-j*, Supplementary Table 3**. We clustered these genes according to their spatially resolved projections using a Bayesian spatial-correlation approach, resulting in 3 different patterns. These spatial patterns were found to be highly overlapping with our prior clustering, suggesting that cells of the hypoxia-reactive states were spatially congruent to pattern 1, and the immune-reactive states were represented in pattern 2, ***Extended Figure 4e***. The lineage states, predominantly NPC-like and OPC-like, were present in pattern 3, while the radial glia overlapped with reactive and lineage patterns ***Extended Data Figure 5 i-k***. Our findings suggest that the observed intra-tumor heterogeneity involves individual lineages mirroring brain development, which is consistent with the findings of others^3,19^. However, we also observed reactive states in response to various pathological conditions reminiscent of transcriptional signatures reported for reactive astrocytes^20–25^.

### Immune or metabolic environment drives spatially exclusive cell fates

To comprehensively interrogate dynamic adaptations, we annotated the RNA-velocity of all tumor cells (InferCNV-analysis: gain Chr7 and loss Chr 10), ***Figure 2c-d***, and realigned all cells according to their lineage origin, presented by the first 3 principal components, ***Figure 2e***. Macro states including initial and terminal states were estimated by Markov chains based on annotated RNA velocity and transcriptomic similarity (CellRank^26^). Our data indicated that reactive states likely arose from former phylogenetic lineage-differentiated origins, ***Figure 2f***. The identified transcriptional programs that drive the reactive transformation were closely related to signature genes of known states of reactive astrocytes^21^ (*CHI3L1, C1S*), ***Figure 2g***. We observed that initial and terminal states undergo dynamic shifts, and the direction of cellular differentiation was not unambiguously determined, reflecting the enormous plasticity of glioblastomas, ***Figure 2h-j, Extend Data Figure 6***. To evaluate the impact of metabolic alterations in the observed reactive patterns, we performed Matrix-assisted Laser Desorption/Ionization Fourier Transform Ion Cyclotron Resonance imaging mass spectrometry (MALDI-FTICR-MSI) of consecutive slices in six patients (spatial transcriptomic blocks) and traced back metabolic alterations in the regions of unique transcriptional states, ***Figure 3a, Supplementary Table 1***. Spatial metabolomic profiling revealed less intra-patient variability compared to the transcriptional data, suggesting that metabolic heterogeneity based on regional imbalances indeed exists across all patients, ***Figure 3b***. We observed regional alterations of fatty acid metabolism and glycolysis overlapping with signature expression of the hypoxic reactive state, ***Figure 3c-e***. Regions with increased glycolysis also showed additional gains on chromosome 7 and a strong enrichment of genes associated with glycolysis, confirming the consistency of our data.

**Figure 3:**
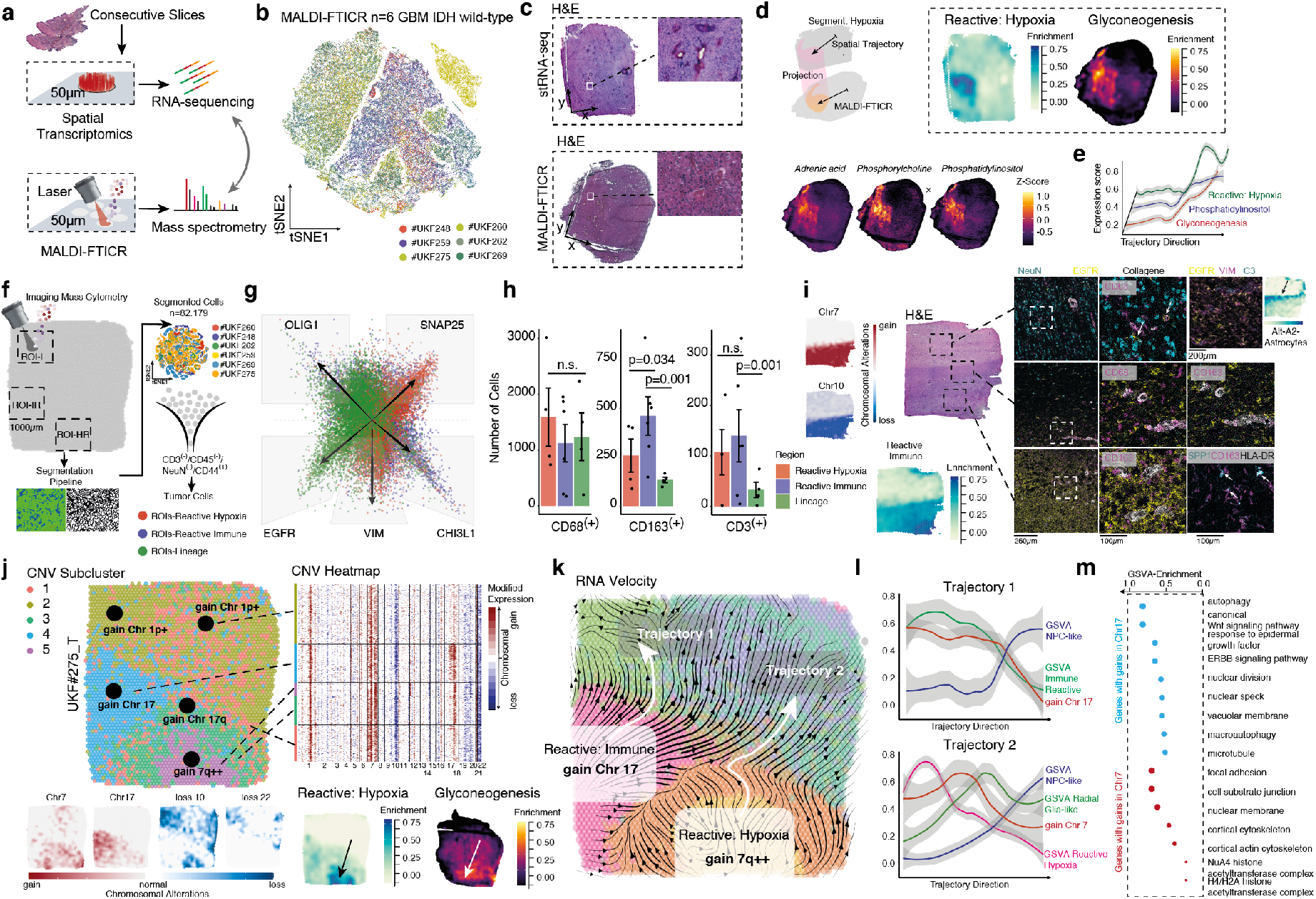
a) Illustration of the workflow. b) Dimensional reduction scatter plot (tSNE) of all batch-corrected specimens (indicated by colors). c) Hematoxylin and eosin stain (H&E) of the spatial transcriptomic sample (upper panel) and the sample for MALDI-FTICR-MSI (bottom panel). Arrows indicate the bottom left side of the sample. d) Spatial overlap of both techniques was performed by manual segmentation (illustrated in the left panel). In the right panel, surface plots of gene set variation analysis or metabolic intensities (z-score enrichment of metabolic pathways) are illustrated. e) Spatial and metabolic intensities are demonstrated along a spatial trajectory (d, upper plot). f) Illustration of the IMC workflow and segmentation pipeline. g) Scatterplot of state-specific markers to determine regional distribution of cell states. Colors indicate the ROIs. h) Bar plots of cell counts in different ROIs. Error bars illustrate the standard error and significance levels were determined by ANOVA. i) Sample with tumor and infiltration areas (#UKF_269). CNV surface plots indicate the chromosomal alterations at spatial resolution (left) indication low tumor penetrance in the upper regions. IMC ROIs are marked in the H&E staining. IMC images (right) from all regions illustrate the different distribution of tumor cells (EGFR), neurons (NeuN) and myeloid cells (CD68 and CD163). Right upper panel, reactive astrocytes (VIM/C3) and GBM cells (EGFR) are presented at the tumor boarder. The enrichment of the alternative-A2-transcriptional signature is illustrated at the right side. Right-bottom, immunostaining (IMC) of SPP1, HLA-DR and CD163 illustrate the typical tumor-associated activated myeloid cells. j) Hierarchical clustering of the estimated CNV alterations is presented (at spatial resolution, right panel) or in a CNV heatmap. At the bottom, CNV surface plots indicate the chromosomal alterations at spatial resolution (left) and the corresponding spatial and metabolic intensities (enrichment). k) RNA-velocity stream at spatial resolution, colors indicate the SNN clusters. Arrows mark spatial trajectories along the velocity streams. l) Line plots of both trajectories demonstrate the gene set enrichment of subtypes and chromosomal alterations along the velocity streams. m) Gene set enrichment analysis of the 25% most altered genes (estimated CNV score) on chromosome 7 and 17.

To understand and validate our findings at single-cell resolution, we performed imaging mass cytometry of consecutive sections (6 patients, 14 different 1000µm regions of interest, ROI) resulting in a comprehensive proteomic map of 82.179 cells after segmentation, **Figure 3f, *Supplementary Table 1***. Based on state specific markers, we confirmed the distribution of GBM cells within ROIs of lineage, reactive immune or hypoxia differentiation, **Figure 3g**. In particular, we found that T cells CD3(+) were preferentially localized in regions of tumor cells with a reactive differentiation without significant differences between hypoxic and immune reactive regions. CD68(+) myeloid cells were similarly distributed across all reactive- and lineage-state-ROIs, however, CD163(+) myeloid cells were significantly enriched in immune reactive ROIs (ANOVA, p=0.001), **Figure 3h**. By mapping the different spatial levels of tumor infiltration, we found that activated myeloid cells marked by CD163, SPP1 and HLA-DR were enriched in regions of reactive inflammation GBM state, **Figure 3i**. Additionally, we showed that GBM cells and GBM-associated reactive astrocytes VIM(+)/C3(+) form a scar-like formation at the tumor border.

### Spatiotemporal lineages and transcriptional plasticity in glioblastoma

Based on the assumption that environmental conditions shape transcriptomic states, we aimed to explore to what extent these conditions cause selective pressure, leading to more resistant tumor subclones. Using a hidden Markov model, we predicted the spatially resolved subclonal architecture, ***Figure 3j***. We found that only a subset of patients revealed subclones as defined by different CNVs in our examined regions. These patients showed a non/small overlap between individual subclones, leading to the assumption that the subclonal architecture was not randomly distributed, ***Figure 3k***. We estimated the pseudotemporal hierarchy using RNA velocity, which demonstrated a large variance of bidirectional subtype shifts across subclonal regions, and highlights the transcriptional plasticity of GBM’s, ***Extended Data Figure 7***. A less common alteration of chromosome 17 was correlated with the enrichment scores of the reactive immune subclass, ***Figure 3j-k and Extended Data Figure 7***. The upper 0.25 quantile of altered genes on chromosome 17 showed a pathway enrichment in Wnt/β-catenin (Wnt), which is known to subvert cancer immunosurveillance^27^, and in ErbB protein family signaling, **Figure 3m**. A spatial overlap of gains in chromosome 7 and hypoxic-related signature enrichment was observed, which followed the same pattern along the RNA-velocity stream (Trajectory 2), ***Figure 3k-m***. Enrichment analysis of the most altered genes on chromosome 7 revealed dysregulation of focal adhesion and of the actin cytoskeleton, suggesting increased migratory capacity which may be required for escape from metabolic imbalance, **Figure 3m**.

### Patient-specific spatially resolved gene expression is driven by age

Our analysis revealed that environmental factors shape distinct transcriptional programs, which partially explained the high inter-patient variance. Global changes of the neural environment which arise during aging remain less explored but are of high importance. Several neurological diseases such as Alzheimer’s disease (AD) or Multiple Sclerosis (MS) cause a general inflammatory environment and drive the inflammatory transformation of glia cells^17,28,29^. An increase of inflammatory transformation was also reported for the aging brain, which could be caused by damage to the blood-brain-barrier^30^. We hypothesized that age-related alterations in the neural environment may also support glioblastoma transcriptional plasticity and differentiation. Indeed, we observed an unbalanced age distribution within our identified transcriptional subclasses which revealed a shift towards increased reactive adaptation within elderly patients, ***Figure 4a***. In order to elucidate the biological significance of aging in GBM and in the human brain, we acquired spatial transcriptomic datasets (n=6) from non-malignant specimens across different age groups, ranging from 19 to 81 years. We confirmed the absence of malignant cells by inferring somatic alterations, ***Extended Data Figure 8a-b***. Common markers of reactive astrocytes (*GFAP, CHI3L1* and *C1R*) were up-regulated in elderly cortical specimens, ***Figure 4c***, leading to the assumption that the aging environment may also contribute to the reactive transformation seen in GBM. To underpin our hypothesis, we estimated common age-related gene expression meta-modules and identified genes which were associated with neural differentiation and plasticity (*ENC1, SNAP25 VSNL1*), all of which were significantly downregulated in elderly patients ***Figure 4d and Extended Data Figure 8c-f***. Common markers of reactive astrocytes such as *GFAP, CHI3L1* and oligodendrocytes (*MBP, PLP1*) were upregulated, ***Figure 4e and Extended Data Figure 8h***. Through integration of age-related co-expression modules from cortex and tumor samples, we identified a shared inflammatory activation along the estimated temporal trajectory, ***Figure 4f***. This corroborated our assumption that the age-related alterations of the neural environment shapes heterogeneity and cellular differentiation in GBM which was further confirmed by weighted correlation network analysis using bulk RNA-seq analysis, ***Extended Figure 9***.

**Figure 4:**
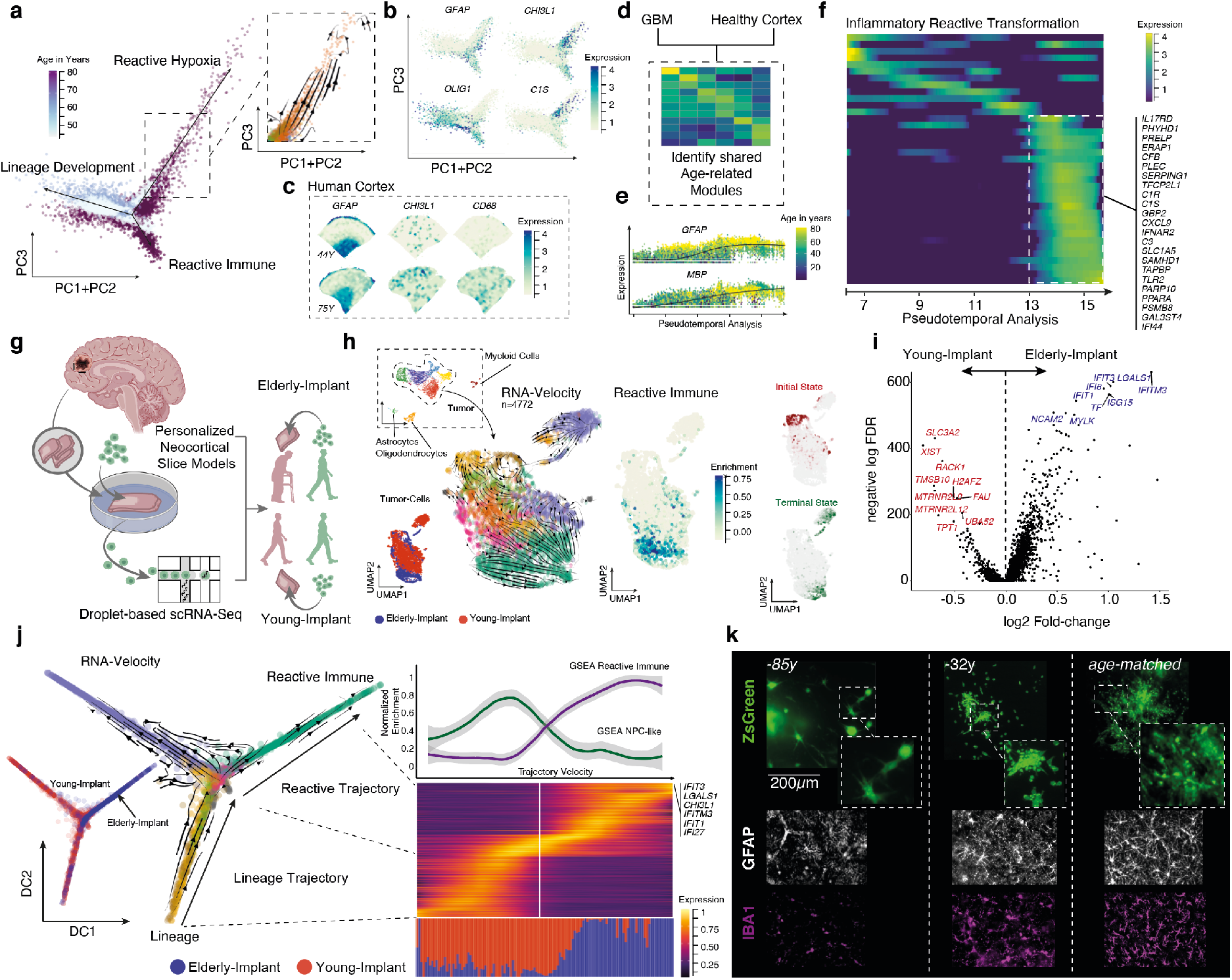
a) Dimensional reduction (see Fig.2 for detailed explanation) of the first three eigenvectors, colors indicate the age of patients. Predominantly, highly dynamic branches (right panel) mostly occupied by elderly patients. b) Dimensional reduction with gene expression of inflammatory/reactive astrocyte genes. c) Surface plots of non-neoplastic cortex sample of a young (upper row) and an elderly patient (lower row). d) GBM and non-neoplastic samples are used to estimate age-related gene expression modules. e) Scatter plot of pseudotemporal depended gene expression, colors indicate patient age. f) Heatmap of gene expression along the estimated pseudotemporal axis. g) Illustration of the neocortical slice model. h) Dimensional reduction (UMAP) of scRNA-seq. UMAPs indicate different sample source (left bottom), RNA-velocity (middle panel) and enrichment of reactive/lineage marker expression (right plots). The estimated initial and terminal states are illustrated on the right. i) Volcano plot of differential gene expression of tumor cells injected into young- or elderly brain slices. j) Diffusion plots (dimensional reduction, with RNA-velocity) indicate the difference of reactive and lineage differentiation along the major axis. Heatmaps of the lineage to reactive trajectory are illustrated. On the bottom of the heatmap, a barplot indicates the sample source. A lineplot at the top of the heatmap illustrates the gene set enrichment analysis of the reactive immune and NPC-like cell states. k) Immunostainings of an elderly tumor which was injected into young (left), middle-aged(middle) and age-matched (right) cortical slices. Immunostainings of GFAP and IBA1 indicate (bottom plots) the increased number of reactive glia during aging.

### Age influences GBM growth and heterogeneity

Of note, in addition to the investigated tumor- and age-related signaling alterations, GBM commonly occurs between the 6th and 8th decade of life, implying that the ageing environment contributes to malignant transformation. To experimentally validate this hypothesis, we used an advancement of our most recently described novel human neocortical slice model because age-related impacts are difficult to investigate in murine models. We injected a patient-derived, Zs-green tagged GBM cell line (38y) into cortical slices from a young (15y, n=6) and an elderly donor (63y, n=7) ***Figure 4g***. After 7 days of culture, we FACS sorted tumor cells and performed scRNA-seq. A total of 5672 cells were obtained, from which 4772 were identified as tumor cells based on their CNV alterations, ***Figure 4i, Extended Data Figure 10a-e***. Tumor cells injected into the elderly cortical slices revealed lower heterogeneity and a strong enrichment of reactive markers, ***Figure 4i***. Using pseudo-temporal reconstruction of RNA-velocity and cell fate determination (CellRank), we found that terminal states predominantly contained reactively transformed cells ***Figure 4h***. Additionally, tumor cells obtained from aged cortex slices were more abundant in the terminal-reactive population, ***Figure 4h***. Next we performed differential gene expression analysis to decipher the impact of an aged-environment on tumor cells which revealed an up-regulation of genes associated with INF gamma response (*IFIT3, IFI6* and *IFIT1*), ***Extended Data Figure 10f***, and a loss of known markers of development programs (*H2AFZ also referred to as H2AZ1*)^2^. Diffusion map re-embedding of the scRNA-seq data indicated a major branching for developmental and reactive programs. Along a trajectory ranging from development to reactive cell fates, we observed an increasing accumulation of tumor cells from aged cortex slices, ***Figure 4j***.

To further validate the impact of age-related microenvironmental alterations on growth behavior of GBM’s, we injected GBM cells derived from both elderly and young patients into cortical slices from a variety of age groups including infantile, middle aged and elderly donors, resulting in significantly reduced growth rate compared to the younger cell line, ***Figure 4k and Extended Data Figure 10g***. Optimal tumor growth was obtained in age-matched slices, ***Extended Data Figure 10g***. Here, we present a novel approach to illuminate the increased incidence and poorer prognosis of glioblastoma in the elderly. These insights also affect the further design of tumor models, as so far little attention has been paid to age-related effects.

## Conclusion

Our investigation uncovered novel insights into the bi- and unidirectional interactions between microenvironment and transcriptional heterogeneity across time and space in glioblastoma. The in-depth, spatially resolved characterization of glioblastoma at various molecular levels facilitates the discovery of the dynamic adaptation of cellular states and spatial relationships within the tumor microenvironment. In close proximity to developmental trajectories of the brain or adaptive transformation in various CNS diseases, we uncovered dynamic differentiation of GBM cells along lineage developmental states and reactive transformations. Deciphering the pathogenesis of each state demonstrated a close link between metabolic alterations and inflammatory responses as drivers of reactive adaptation in GBM cells. We demonstrated that age-induced inflammatory processes are the major cause of transcriptional shift towards reactive states in elderly GBM patients. Using our human neocortical GBM model across different age groups, we confirmed that glioblastoma heterogeneity and plasticity is age-related. This suggests that artificial age differences in tumors models lead to spurious experimental results regarding tumor growth and transcriptional plasticity. Our results suggest that glioblastoma adapts to the aging brain, necessitating tailored therapeutic approaches and underpinning the importance of a personalized approach in neuro-oncology.

## Methods

### Ethical Approval

The local ethics committee of the University of Freiburg approved the data evaluation, imaging procedures and experimental design (protocol 100020/09 and 472/15_160880). The methods were carried out in accordance with the approved guidelines, with written informed consent obtained from all subjects. The studies were approved by an institutional review board. Further information and requests for resources, raw data and reagents should be directed and will be fulfilled by the Contact: D. H. Heiland. A complete table of all materials used is given in the supplementary information.

### Spatial Transcriptomics

The spatial transcriptomics experiments were performed using the 10X Visium Spatial Gene Expression kit (https://www.10xgenomics.com/spatial-gene-expression). All the instructions for Tissue Optimization and Library preparation were followed according to manufacturer’s protocol. Here, we briefly describe the methods followed using the library preparation protocol.

### Tissue collection and RNA quality control

Fresh tissue collected immediately post resection was quickly embedded in Tissue-Tek^®^ O.C.T.^™^ Compound (Sakura, 4583) and snap frozen in isopentane pre-chilled in liquid nitrogen. Embedded tissue was stored at −80°C until further processing. A total of 10 sections (10μm each) per sample were lysed using TriZOl (Invitrogen, 15596026) and used to determine RNA integrity. Total RNA was extracted using PicoPure RNA Isolation Kit (Thermo Fisher, KIT0204) according to the manufacturer’s protocol. RIN values were determined using a Fragment Analyzer 5200 (RNA kit, Agilent, DNF-471) according to the manufacturer’s protocol. It is recommended to only use samples with an RNA integrity value >7.

### Spatial Gene Expression Protocol

10 µm thick sections were mounted onto spatially barcoded glass slides with poly-T reverse transcription primers, with one section per array. Slides were fixed in 100% methanol and H&E staining was performed. Brightfield imaging was done at 10x magnification with a Zeiss Axio Imager 2 Microscope, and post-processing was performed using ImageJ software. Following imaging, permeabilization took place for a pre-determined time to release and capture mRNA from the tissue onto primers on the slide. Template switch oligos were introduced in order to generate a second strand in a reverse transcription reaction and produced second strand was cleaved off by denaturation. Next, generated cDNA was amplified and fragments in the size of interest were selected using SPRIselect reagent (Beckman Coulter, B23318). Quality check was performed using a Fragment Analyzer (HS NGS Fragment kit, Agilent, DNF-474). Further, fragmentation and double-sided size selection using SPRIselect reagent was carried out in order to optimize cDNA fragments for Illumina NextSeq Sequencing System. Unique indexes as well as P5 and P7 Illumina primers were added to the libraries. The average length of the final libraries was quantified using a Fragment Analyzer (HS NGS Fragment kit, Agilent, DNF-474) and the concentration of libraries was determined using a Qubit 1X dsDNA HS kit (Thermo Fisher, Q33231). Final libraries were diluted to 4nM, pooled and denatured before sequencing on the Illumina NextSeq 550 platform using paired-end sequencing. We used 28 cycles for read 1, 10 cycles per index and 120 cycles for read 2 on a NextSeq 500/550 High Output Kit v2.5 (Illumina, 20024907).

### Data Import and preprocessing, filtering and normalization

Data were analyzed and quality controlled by the cell ranger pipeline provided by 10X. For further analysis we developed a framework for spatial data analysis. The cell ranger output can be imported into SPATA by either a direct import function (SPATA:: initiateSpataObject_10X) or manually imported using count matrix and barcode-coordinate matrix as well the H&E staining. The routine import applies following steps via the Seuratv4.0 package: To normalize gene expression, values of each spot were divided by the estimated total number of transcripts and multiplied by 10,000, followed by natural-log transformation. As described for scRNA sequencing, we removed batch effects and scaled data by a regression model including sample batch and percentage of ribosomal and mitochondrial gene expression.

### Dimensional reduction

We used the 2000 most variable expressed genes and decomposed eigenvalue frequencies of the first 30 principal components. We used either the PCA analysis implemented in Seuratv4.0^31^ or a generalized principal component analysis (GLM-PCA) for non-normal distributions^32^ due to the fact that our UMI counts follow multinomial sampling with no zero inflation. The obtained components were used for shared nearest neighbor-Louvain (SNN-Louvain) clustering followed by nonlinear dimensional reduction using the UMAP or tSNE algorithm. We estimated diffusion maps by the destiny package^33^.

### Clustering and benchmarking

For all cluster approaches of spatial transcriptomics and single-cell RNA-seq we used the non-trivial estimated eigenvectors. An euclidean distance matrix was computed to identify pairs of cells with shared neighbors similar to the SNN-Cliq approach^34^. Cluster integrity was estimated by the highest modularity of each cluster from a graph, based on random connections between nodes^35^. Additionally, we benchmarked our results by hierarchical clustering, k-Means and Partitioning Around Medoids in which the optimal k was estimated by gap-statistics. Classical cluster comparison was not performed on the full dataset due to memory constrains. Cluster with less than 100 spots or less than 20 significantly differently expressed genes were excluded or defined as outliers. Estimation of the cluster marker genes was performed by the SPATA implementation of a Wilcoxon sum-rank test.

### Identification of shared transcriptional programs and gene expression modules across patients

First, we performed cluster analysis (SNN, as described above) of malignant spots from each tumor separately. Selection of meaningful clusters was performed as described above and benchmarked by various cluster approaches. For each individual cluster, we estimated the number of significantly expressed genes by the following criteria: Genes with 2.5-fold increase of the average log fold-change and corresponding p values below 0.05 (False-Discovery Rate of a Wilcoxon Rank Sum test). In order to ensure non overlapping individual clusters, we merged clusters with a Jaccard index above 70%. Genes of each clusters were used as cluster signatures for further processing. In the next step, we estimated the cluster similarity using Jaccard indices and discarded clusters with a lower index than 0.2. Next, we extracted genes with were represented in more than 70% of all clusters to identify common expressed signature genes. Using hierarchical clustering of the signature genes by average linkage, we identified six modules containing 309 genes. We performed benchmarking of our clustering by k-Means and Partitioning Around Medoids in which the optimal k was estimated by gap-statistics.

### Pattern recognition and clustering

First, we sought for spatially exclusive expressed genes also referred to as spatial expression (SE) using a generalized linear spatial model implemented in the SPARK algorithm^36^. Through this approach we analyzed each tumor separately and selected all significant SE genes (threshold p corrected by Benjamini–Hochberg p<0.001). For further spatial pattern analysis, we selected genes which were present in at least 75% of all tumors. To unravel the spatial arrangement and detect co-localized patterns, we estimated spatial co-localization by a Bayesian spatial correlation model of all recurrent SE genes. This resulted in a correlation matrix which was hierarchically clustered and revealed 5 distinct patterns. We further summarized these patterns into three major modules based on our findings from our first approach. The two reactive patterns (hypoxia and immune-related genes) showed distinct from each other while developmental subcluster (OPC and NPC) revealed a stringer overlap.

### Pathway analysis of gene sets

We performed pathway analysis by three different methods all implemented into our SPATA toolbox. As presented in our figured we used gene set variation analysis (GSVA) or z-scored enrichment of gene sets. The analysis was performed through the GSVA package^37^. For GO-term enrichment we used the DOSE package and cluster profiler^38^.

### Comparison of cortex and tumor samples

First, we merged all cortex samples (n=5) with a total number of 17.275 transcriptomes. For batch effect removal, we read the data into a monocle3^39^ object and aligned samples by matching mutual nearest neighbors (monocle3::align_cds)^40^. Next, we performed pseudotime analysis by setting the root into spots from a 19-years old cortex sample. The estimated mean pseudotime per sample and real age showed a significant correlation (R^2^=0.56 p<0.031). To detect genes which are differentially expressed along our estimated age-trajectory, we performed Moran’s I statistics, a measure of multi-directional and multi-dimensional spatial autocorrelation^41^. We merged genes into modules which were co-expressed across all spots using the monocle3:: find_gene_modules() function. Next, we performed similar steps using tumor samples (with altered CNVs) and compared modules by similarity using the Jaccard-index. We identified a shared module which was highly enriched in elderly patients containing immune related gene expression.

### Weighted correlation analysis of the TCGA database

In order to confirm the increase of inflammatory genes in elderly patients we performed a weighted correlation network analysis (WGCNA) with age as a co-variable^42^. The TCGA gene expression dataset (RNA-seq Bulk GBM) was downloaded from the GlioVis database^43^. In a first step, we estimated the soft-thresholding power (sft) which was required to reach scale-free topology by iterating over *p= 1…, 10*. Using an unsigned network architecture, we reached scale-free topology at a sft of 5. We performed block wise WGCNA using a Pearson-correlation measurement and a deep split of 2. Next, we merged modules with highly correlating eigengenes (WGCNA:: mergeCloseModules) and estimated the eigengene-based connectivity (kME). We correlated the age of patients and the identified kME which revealed a significant correlation to the kMEmagenta. Next, we characterized the significant correlation modues by GO-term enrichment analysis and confirmed the inflammatory activation in elderly patients.

### RNA velocity estimation

We used the CellRanger BAM file to separate expression matrices of spliced and unspliced reads through the ready-to-use pipeline from the velocyto package^44^. The resulting .loom file was read into the scVelo Seurat wrapper (https://github.com/satijalab/seurat-wrappers). We merged the Seurat objects and performed batch effect removal as explained above. After data integration, Seurat objects with exonic and intronic gene-level UMI counts were converted into h5ad format (https://github.com/mojaveazure/seurat-disk). We read-in the h5ad files to an AnnData object. Next we performed normalization and selected the 2,000 most variable expressed genes by the scVelo package (v0.2.3)^45^. We excluded all genes with less than 20 assigned reads across the exonic and intronic components and estimated RNA velocity and latent time using the dynamical model. Data will be exported as .csv files and implemented into a SPATA object for further visualization. The explained pipeline is implemented into a SPATA wrapper for scVelo (SPATA::getRNA velocity, in the development branch).

### Infer lineage differentiation by CellRank

After performing the dynamical model, we estimated macro states which represent initial, terminal states as well as transient intermediate states using the CellRank package (v1.1.0, https://github.com/-theislab/cellrank)^26,45^. We constructed a transition matrix using the connectivity kernel which was analyzed by Generalized Perron Cluster Cluster Analysis (GPCCA)^46^ after computing a Schur triangulation. We estimated the probability of all identified macro state (initial and terminal states) in each spot. The probability vectors are implemented into the fdata slot of the corresponding SPATA object. Lineage driver genes of each estimated macrostate were identified by the *compute_lineage_drivers* function of CellRank. Additionally, we used the partition-based graph abstraction (PAGA) to simplify state transition in space.

### Visualization of RNA velocity in spatial transcriptomic datasets

Visualization off all tumor samples was performed by using the first 3 principal components (PC1-3) which was integrated into the AnnData object in the adata.obsm[‘X_umap’] slot. The velocity streams were computed by the pl.velocity_embedding_stream function referring to the “X_umap” slot. In our spatial transcriptomic data, we aimed to preserve the spatial architecture when adding the velocity streams. We migrated the spatial coordinates from the SPATA object to the AnnData object into the adata.obsm[‘X_umap’] slot which was used for the pl.velocity_embedding_stream function.

### Estimation of transient gene expression programs along RNA velocity streams

In order to estimate transcriptional programs which were dynamically regulated in space (spatial transcriptomics) and time (RNA velocity estimation) we used the computed velocity streams as spatial trajectories. Using the *SPATA::createTrajectories* function, we sought for genes which followed a predefined dynamic along our spatio-temporal trajectory as recently described^47^.

### Spatial gene expression

The visualization of spatial gene expression is implemented in the SPATA software SPATA:: plotSurfaceInteractive. For spatial expression plots, we used either normalized and scaled gene expression values (to plot single genes) or scores of a set of genes, using the 0.5 quantile of a probability distribution fitting. The x-axis and y-axis coordinates are given by the input file based on the localization at the H&E staining. We computed a matrix based on the maximum and minimum extension of the spots used (32×33) containing the gene expression or computed scores. Spots without tissue covering were set to zero. Next, we transformed the matrix, using the squared distance between two points divided by a given threshold, implemented in the fields package (R-software) and adapted the input values by increasing the contrast between uncovered spots. The data are illustrated as surface plots (plotly package R-software) or as images (graphics package R-software).

### Spatial correlation analysis

In order to map spatial correlated gene expression or gene set enrichments we used z-scored ranked normalized expression values. One gene expression vector or enrichment vector of a gene set is used to order the spots along a spatial trajectory. We construct the trajectory of spots from lowest ranked to highest ranked spot (based on z-scored input vectors). The genes of interest (which were correlated with the spatial trajectory) are fitted by loess-fit from the stats-package (R-software) and aligned to the ranked spots and scaled. Correlation analysis was performed by Pearson’s product moment correlation coefficient. For heatmap illustration the gene order was computed by ordering the maximal peak of the loess fitted expression along the predefined spatial trajectory.

### Identification of cycling cells

We used the set of genes published by Neftel and colleagues^1^ to calculate proliferation scores based on the GSVA package implemented in R-software. The analysis based on a non-parametric unsupervised approach, which transformed a classic gene matrix (gene-by-sample) into a gene set by sample matrix resulted in an enrichment score for each sample and pathway. From the output enrichment scores we set a threshold based on distribution fitting to define cycling cells.

### CNV estimation

For CNV analysis we implemented a CNV pipeline into our SPATA R tool available in the development branch, https://github.com/theMILOlab/SPATA. Copy number Variations (CNVs) were estimated by aligning genes to their chromosomal location and applying a moving average to the relative expression values, with a sliding window of 100 genes within each chromosome, as described recently^8^. First, we arranged genes in accordance to their respective genomic localization using the InferCNV package (R-software)^8^. As a reference set of non-malignant cells, we used a spatial transcriptomic dataset from a non-malignant cortex sample. To increase speed and computational power, a down-sampling is optional possible. To avoid the considerable impact of any particular gene on the moving average we limited the relative expression values [-2.6,2.6] by replacing all values above/below *exp(i)*=|2.6|,by using the infercnv package (R-software). This was performed only in the context of CNV estimation as previous reported^48^. The exported .RDS files were reimported and grouped by chromosomal averages of estimated CNV alterations and aligned to their spatial position using the *fdata* slot of the SPATA object. Using the *SPATA::joinWithFeatures()* function extraction of cluster-wise comparison are performed. Additionally, we implemented the option to select the most altered genes of chromosomes.

### MALDI-FTICR-MSI

Tissue preparation steps for MALDI imaging mass spectrometry (MALDI-MSI) analysis was performed as previously described^49,50^. Frozen tissues were cryo sectioned at 10 µm from the same tissue block as used for spatial transcriptomics and thaw mounted onto indium-tin-oxide coated conductive slides (Bruker Daltonik, Bremen, Germany). The matrix solution consisted of 10 mg/ml 9-aminoacridine hydrochloride monohydrate (9-AA) (Sigma-Aldrich, Germany) in water/methanol 30:70 (v/v). SunCollectTM automatic sprayer (Sunchrom, Friedrichsdorf, Germany) was used for matrix application. The MALDI-MSI measurement was performed on a Bruker Solarix 7T FT-ICR-MS (Bruker Daltonik, Bremen, Germany) in negative ion mode using 100 laser shots at a frequency of 1000 Hz. The MALDI-MSI data were acquired over a mass range of m/z 75-1000 with 50 μm lateral resolution. Following the MALDI imaging experiments, the tissue sections were stained with hematoxylin and eosin (H&E) and scanned with an AxioScan.Z1 digital slide scanner (Zeiss, Jena, Germany) equipped with a 20x magnification objective. After the MALDI-MSI measurement, the acquired data underwent spectra processing in FlexImaging v. 5.0 (Bruker Daltonics, Germany) and SCiLS Lab v. 2020 (Bruker Daltonik GmbH). MS peak annotation was performed using Human Metabolome Database (HMDB, https://www.hmdb.ca/)^51^ and METASPACE (https://metaspace2020.eu/)^52^.

### MALDI data analysis

We read-in the files into R using the *readImzML* function from the cardinal package^53^. We reshaped the pixel data matrix into an intensity matrix and a matrix of coordinates for each tumor separately. We filtered the m/z matrix to annotated peaks (METASPACE database) using the *match*.*closest* function from the MALDIquant package resulting in a metabolic intensity matrix^54^. The intensity matrix and the corresponding spatial coordinated were imported into a SPATA object for further spatial data analysis using the SPATA::initiateSpataObject_MALDI.

### Human Organotypic Slice Culture

Human neocortical slices were prepared as recently described^21,55^. Resected cortical tissue (assessed by EEG and MRI) was immediately brought to the lab in the “preparation medium” (Gibco Hibernate™ media supplemented with 1 mM Gibco GlutaMax™, 13 mM Glucose, 30 mM NMDG and 1% Anti-Anti) saturated with carbogen (95% O2 and 5% CO2). Capillaries and damaged tissue were dissected away from the tissue block. The combo of GlutaMax and NMDG in the collection medium has provided us with best tissue recovery post resection. 300 μm thick cortical slices were obtained using a vibratome (VT1200, Leica Germany) and incubated in preparation medium for 10 minutes before plating to avoid any variability due to tissue trauma. Tissue blocks (1 cm × 2 cm) typically permits preparation of 18–20 sections. One to three sections were gathered per insert, with care to prevent them from touching each other. The transfer of the slices was facilitated by a polished wide mouth glass pipette. Slice were maintained in growth medium containing Neurobasal L-Glutamine (Lot No. 1984948; Gibco) supplemented with 2% serum-free B-27 (Lot No. 175040001; Gibco), 2% Anti-Anti (Lot No. 15240-062; Gibco), 13 mM d-glucose (Lot No. RNBG7039; Sigma-Aldrich), 1 mM MgSO4 (M3409; Sigma-Aldrich), 15 mM Hepes (H0887; Sigma-Aldrich), and 2 mM GlutaMAX (Lot No. 1978435; Gibco) The entire medium was replaced with fresh medium 24 hours post plating and every 48 hours thereafter.

### Human ex-vivo Glioblastoma Model

ZsGreen tagged BTSC#233 and BTSC#168 cell lines were cultured and prepared as described previously^21^. Briefly, post trypsinization, a centrifugation step was performed, following which the cells were harvested and re-suspended in PBS for 20,000 cells/µl. Cells were then used immediately for injection onto tissue sections. A 10 µL Hamilton syringe was used to inject 1 µL of GBM cells onto the white matter part of the section. Sections with injected cells were incubated at 37°C for a week and culture medium was refreshed every alternative days. Tumor proliferation was monitored by regular fluorescence imaging by means of an inverted microscope (Observer D.1; Zeiss). After a week, sections were either fixed and used for immunostaining or for single cell sequencing.

### Single cell suspension from cultured slices

Nine sections per condition were processed using C-Tubes (Miltenyi Biotech, 130-093-237) with a shortened protocol for the Neural Tissue Dissociation Kit (T) (Milteny Biotech, 130-093-231). Briefly, the tissue as well as the first enzyme mix, containing enzyme T and buffer X, were transferred to a C-tube and incubated at 37°C for 5 minutes, followed by a rotation for 2 minutes. Next, second enzyme mix, containing enzyme A and buffer Y, was added and incubated for 5 minutes, followed by another rotation for 2 minutes. The sample was then filtered and centrifuged in a 50ml falcon and cell pellet was further used for cell sorting.

### Cell sorting for scRNA-seq

Freshly prepared cell suspensions were washed with FACS buffer containing 2% FCS and 1mM EDTA in PBS and stained with DAPI. Cells were sorted on the BD FACSAria™ Fusion flow cytometer at the core facility, University of Freiburg. To gather viable tumor cells, Zs-green positive, DAPI negative populations were collected in BSA-coated tubes containing 2% FCS in PBS and prepared for later droplet-based single cell RNA-Sequencing.

### Single cell RNA-sequencing

Single cell RNA-sequencing was performed according to the Chromium Next GEM Single Cell 3’v3.1 protocol (10x Genomics), based on a droplet scRNA-sequencing approach. In brief, collected cells were added to a prepared master mix containing reagents for a reverse transcription reaction and loaded onto separate lanes of a Chromium Next GEM Chip G. After running the chip on a Chromium Controller, generated GEMs were transferred to a tube strip. Following reverse transcription, GEMs were broken, and cDNA was purified from leftover reagents. Amplified cDNA was fragmented and size-selected using SPRIselect reagent (Beckman Coulter, B23318). i7 indexes as well as P5 and P7 Illumina primers were added to the libraries. The average length of final libraries was quantified using a Fragment Analyzer (HS NGS Fragment kit, Agilent, DNF-474) and the concentration of libraries was determined using a Qubit 1X dsDNA HS kit (Thermo Fisher, Q33231). Final libraries were diluted to 4nM, pooled and denatured before sequencing on an Illumina NextSeq 550 Sequencing System (Illumina, San Diego, CA, USA) using NextSeq 500/550 High Output kit v2.5 (Illumina, 20024906) with 28 cycles for read 1, 8 cycles for i7 index and 56 cycles for read 2.

### Analysis of scRNA-seq

Single cell RNA-seq were processed by 10x Genomics Cell Ranger 3.1.0^56^. Postprocessing was performed by the MILO-pipeline for scRNA-seq (https://github.com/theMILOlab/scPipelines). Single cell analysis was performed by the Seuratv4.0 package and SPATA 1.0 package. We used the Seurat wrapper for scVelo^45^ to performe pseudotime analysis and Cell Rank^26^ for cell fate estimation. After preprocessing of the data through Seurat, we imported the data into SPATA. Further analysis was performed as explained in the sections above.

### Imaging mass cytometry antibody panel

A 39-marker IMC panel was designed including structural and tumor markers as well as markers to assess several innate and adaptive immune cells (**Supplementary Table XX**). Metal-labeled antibodies were either obtained pre-conjugated (Fluidigm) or labeled in-house by conjugating purified antibodies to lanthanide metals using the Maxpar X8 antibody labelling kit (Fluidigm) according to the manufacturer’s instructions. In addition, 89-Yttrium (III) nitrate tetrahydrate (Sigma Aldrich, cat. # 217239-10G) and 157-Gadolinium (III) chloride (Trace Sciences Int.) were diluted in L-buffer to a 1M stock solution and further diluted to a 50 μM working solution for subsequent antibody labelling with the Maxpar X8 labelling kit. Metal-conjugated antibodies were titrated and validated on glioblastoma, brain, liver and tonsil tissue.

### Sample preparation and staining for imaging mass cytometry

10 µm thick tissue sections on SuperFrost plus slides (R. Langenbrinck GmbH, 03-0060) were dried at 37°C for one minute and fixed in 100% methanol for 30 minutes at −20°C. Slides were rinsed three times in TBS for 5 minutes each. Tissue sections were encircled with a PAP pen (ImmEdge, Vector laboratories, H-4000) and blocked for 45 minutes at room temperature using SuperBlock (TBS) Blocking Buffer (ThermoFisher Scientific, 37581). The sections were then stained with a mix of metal-labeled primary antibodies diluted in TBS with 0.5% BSA as well as 10% FBS and incubated at room temperature for one hour. Slides were rinsed in TBS-T (TBS supplemented with 0.2% Tween-20) twice and twice in TBS for 5 minutes each. Tissue sections were then stained with Iridium Cell-ID intercalator (500 μM, Fluidigm, 201192B) diluted 1:2000 in TBS for 30 minutes at room temperature. Slides were rinsed three times for 5 minutes in TBS, dipped in ddH2O for 5 seconds and air-dried. Slides were stored at room temperature until image acquisition.

### Image acquisition

Two to three 1000 μm^2^ images per patient were acquired using a Hyperion Imaging System (Fluidigm). Briefly, tuning of the instrument was performed according to the manufacturer’s instructions. Tissue sections were laser ablated spot-by-spot at 200 Hz resulting in a pixel size/resolution of 1 μm^2^. Preprocessing of the raw data was conducted using the CyTOF software v7.0 (Fluidigm) and image acquisition control was performed using MCD Viewer v1.0.560.6 (Fluidigm).

### IMC data analysis

Raw data were processed by the bodenmiller pipline^57^. For single-cell analysis we segmented the cells based on the nucleus (DNA-staining) using 6 random crops of each image for training. Training was performed by pixel-wise classification using ilastik^58^. We imported the classification trained images into cell profiler to extract single cell intensities of segmented cells. We analyzed the spatially resolved single-cell matrix by SPATA. For import, we used the *SPATA::initiateSpataObject_MALDI()* function and performed batch effect removal between images by matching mutual nearest neighbors^40^. Cluster analysis was performed as explained above.

## Acknowledgements

DHH is funded by the Else Kröner-Fresenius Foundation. The work is part of the MEPHISTO project (PI: DHH and DD), funded by BMBF (Bundes Ministerium für Bildung und Forschung) (project number: 031L0260B). The Deutsche Forschungsgemeinschaft (DFG, German Research Foundation) supports the work of AW (project number: SFB 824 C04). VR, KJ and UGH funded by BMBF (Bundes Ministerium für Bildung und Forschung) (project number: FMT 13GW0230A), We thank Dietmar Pfeifer for here helpful advices. We thank Biorender.com. We thank Stella Maria Carro for her support and the provision of her laboratory facilities and equipment.

## Data availability

Spatial Transcriptomic RNA-Sequencing Data available: (in preparation), Accession codes:… Full scripts of the analysis are available at github, heilandd/Spatia_Transcriptomics, The used software tool is Spatial Transcriptomic Analysis (SPATA) https://github.com/theMILOlab/SPATA and Tutorials at https://themilolab.github.io/SPATA/index.html, sc-RNA-seq analysis are Data available: (in preparation), Accession codes: Analysis tools: VisLabv1.5 https://github.com/heilandd/Vis_Lab1.5, Further information and requests for resources, raw data and reagents should be directed and will be fulfilled by the Contact: D. H. Heiland.

## Extended Data Figures

**Extended Data Figure 1-3:**
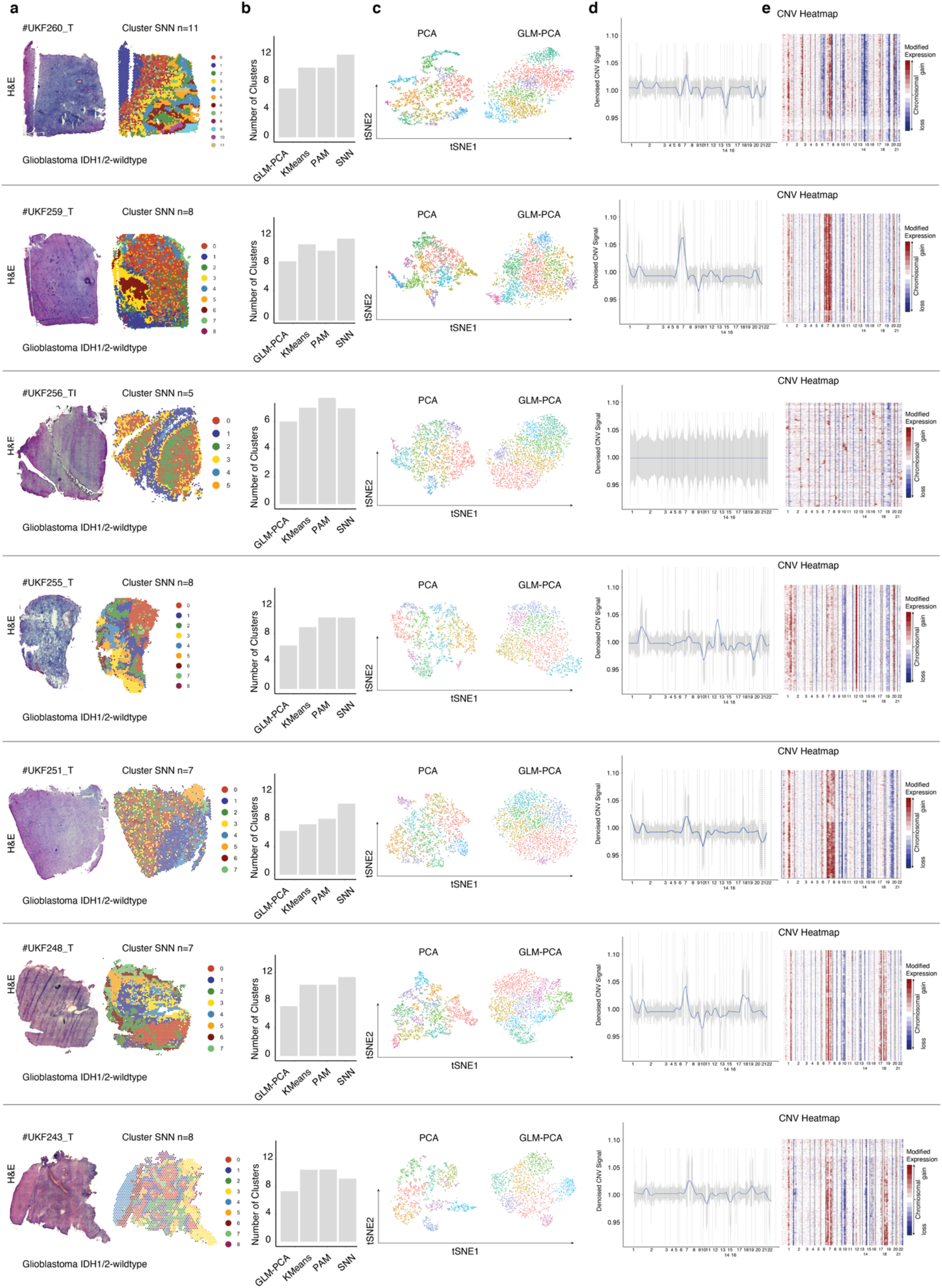
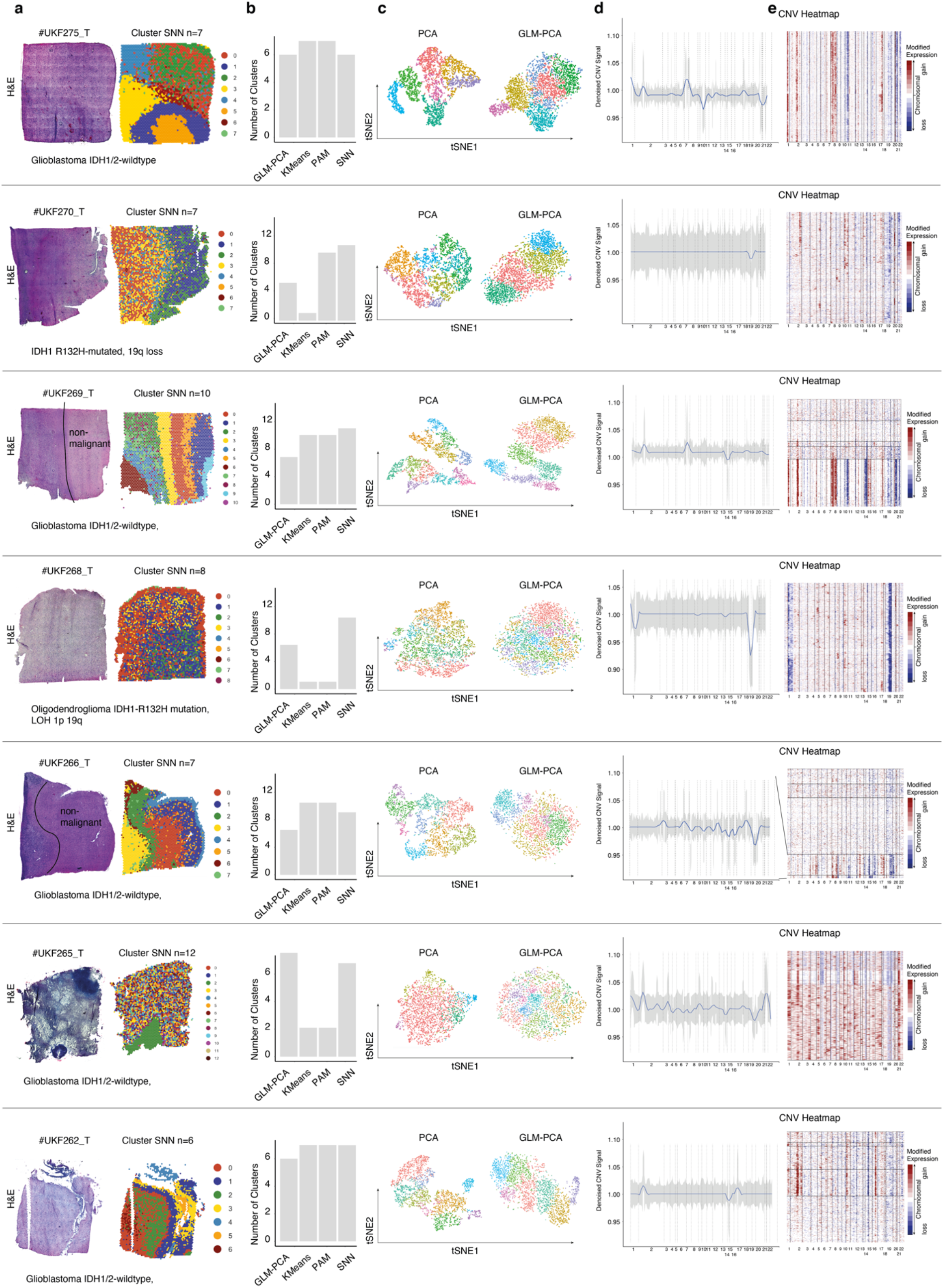
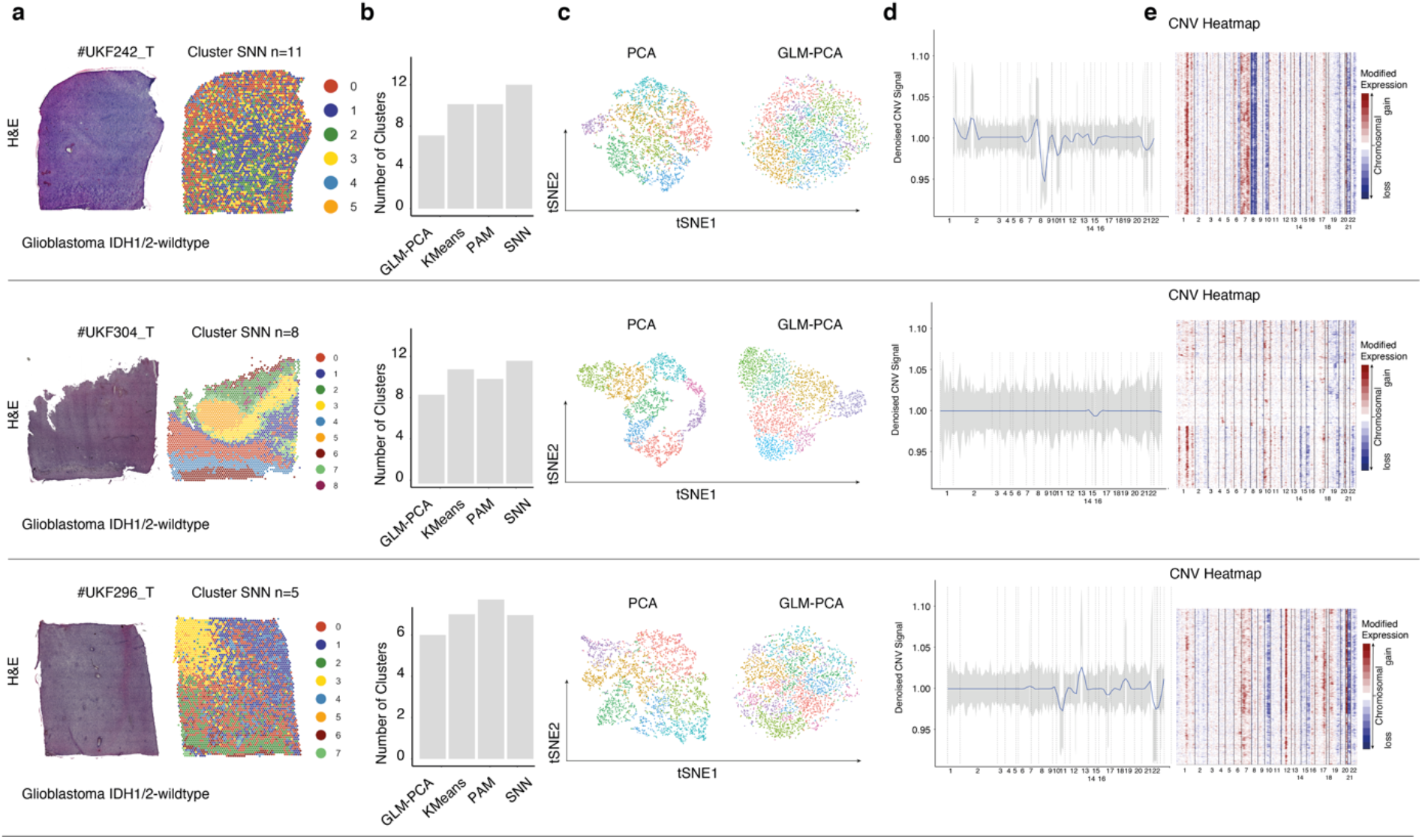
a) Overview of samples with H&E staining (left) and SNN clustering (right) b) Validation of different cluster algorithm. The barplot indicate the estimated optimal number of clusters using a gap statistical approach. c) Dimensional reduction (tSNE) of a classical PCA analysis and a GLM-PCA approach. d) Line plot of sum CNV alterations estimated by InferCNV. The gray area indicates the variance of alterations at each chromosome. e) CNV heatmap with gains in red and losses in blue.

**Extended Data Figure 4:**
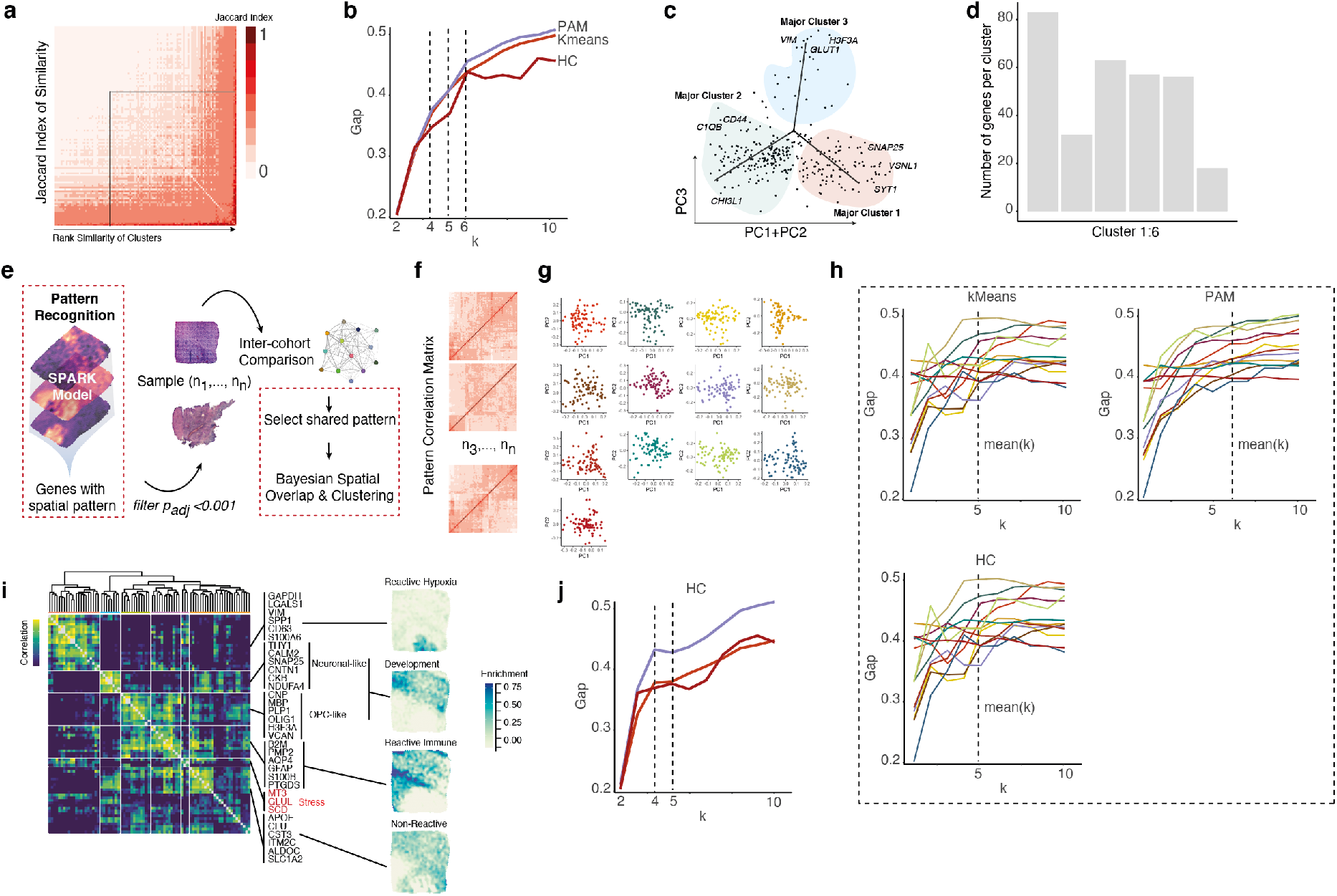
a) Heatmap of shared genes across individual clusters (jaccard index). ∼2/3 of genes are shared across clusters. b) Gap statistic plot of the optimal number of clusters (shared genes of clusters) by various cluster algorithms. i) Dimensional reduction of genes shared in all patients using the first three eigenvectors. c) Number of genes of all identified clusters (signature genes of subclasses). d) Illustration of the pattern recognition approach. e) Example of the distance matrix of genes detected by SPARK. The correspondent PCA plots are illustrated at the right side (f) g) Gap statistics analysis of the optimal number of clusters using different algorithms. Colors indicate the individual patients. h) Heatmap of the three major cluster recognized by hierarchical clustering. i) Gap statistic plot of the optimal number of clusters by various cluster algorithms.

**Extended Data Figure 5:**
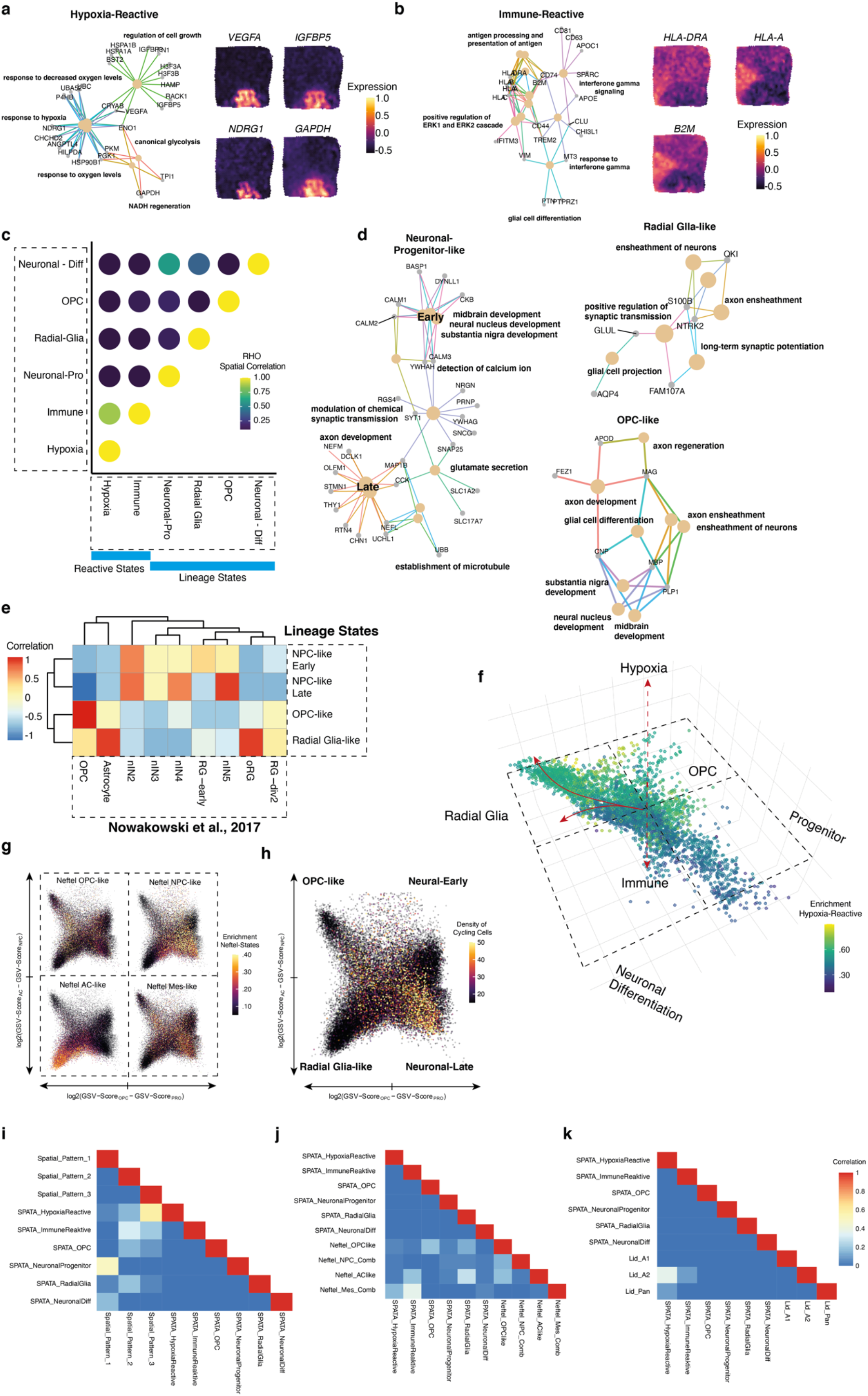
a) Network plot of top enriched pathways of the hypoxic signature. Surface plot of four example genes related to hypoxic response (right) b) Network plot of top enriched pathways of the immune signature. Surface plot of three example genes related to immune response (right) c) Estimated spatial overlap using a Bayesian correlation analysis. The plot indicates signatures occupying similar regions in space. d) Network plot of top enriched pathways of the lineage signatures. e) Analysis of similarity for all lineage stages using the Nowakowski^59^ dataset as reference. f-h) comparison of the Neftel subgroups and the novel signatures. I) Comparison between the signatures of reactive astrocytes, pattern analysis, shared genes approach and the Neftel study.

**Extended Data Figure 6:**
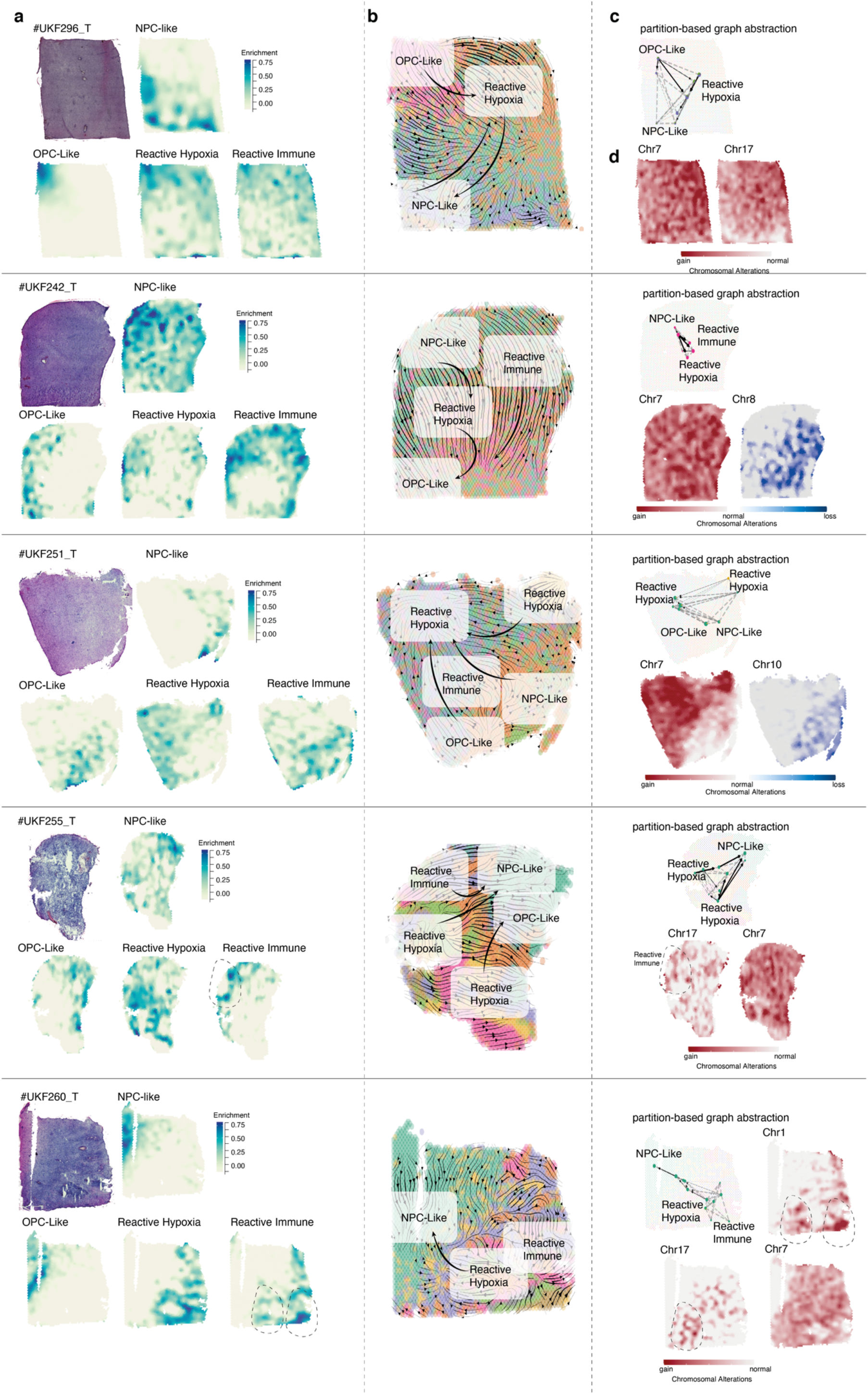
a) H&E staining (left upper) and enrichment surface plots of lineage (NPC- and OPC-like signatures) and reactive genes (hypoxic and immune) b) RNA-velocity stream analysis with arrow which indicate the pseudotemporal development trajectories. Subgroup location is marked as well major differentiation trajectories. c) Aggregation of individual fate maps into a cluster-level fate map using partition-based graph abstraction (PAGA) with directed edges indicates the direction of differentiation at spatial resolution d) Surface plot of estimated CNV alterations of individual patients, red indicate chromosomal gains, blue reveals chromosomal losses.

**Extended Data Figure 7:**
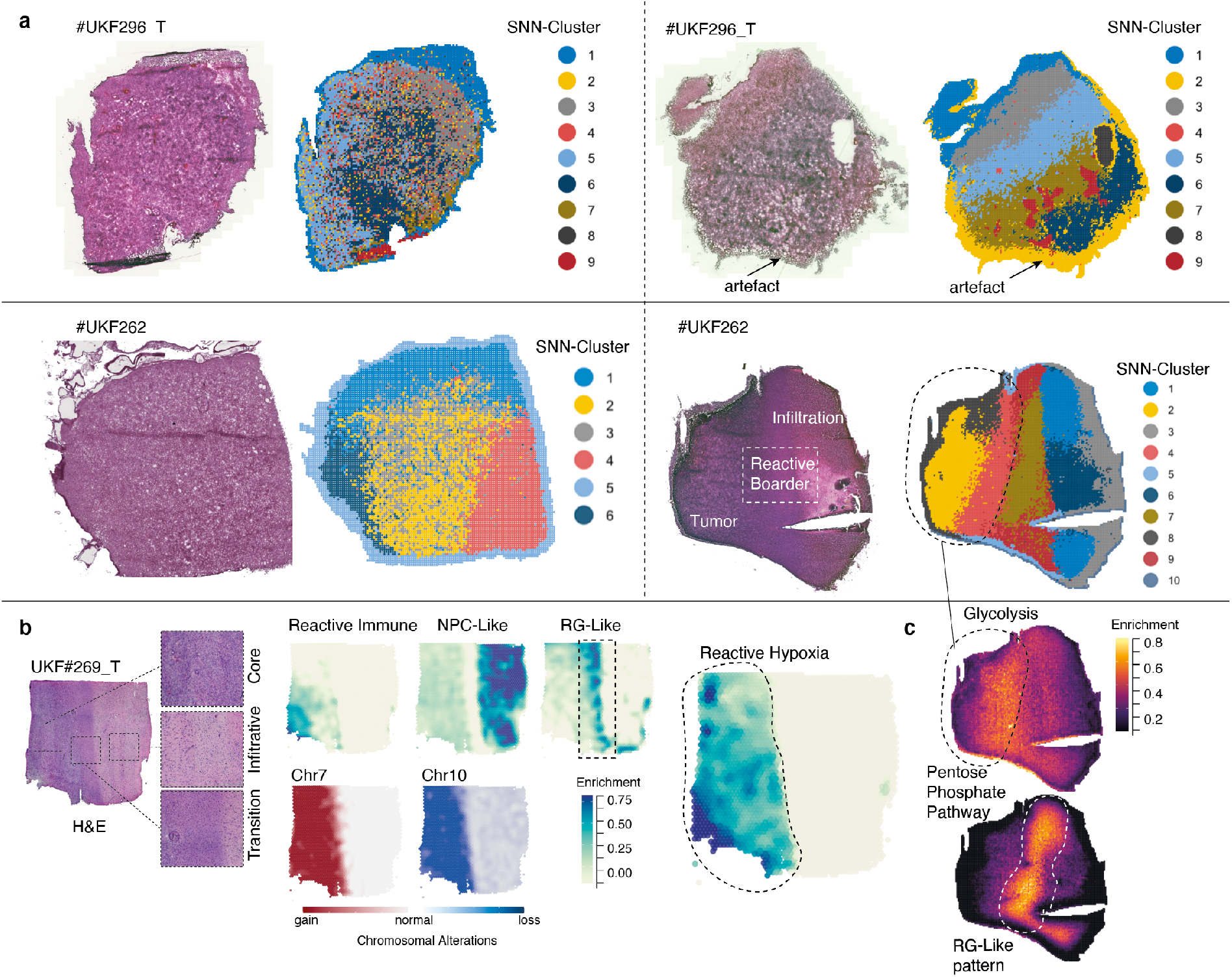
a) H&E staining (left upper) and surface plots with colored clusters (SNN-cluster approach) of MALDI. Clusters with high probability of noisy signal were located at the edge of each sample, most likely indicating a technical artifact. These clusters are excluded for analysis. b) Integration of stRNA-seq and MALDI data indicate regional differences of metabolic processed between tumor core and edge. H&E staining is illustrated at the left side, with magnifications of the three separate areas, namely the tumor core, border or transition area and the infiltrating edge. CNV analysis confirmed the lack of CNV alterations at the infiltrating edge (bottom middle plot). Subtype signatures indicate the enrichment of hypoxic areas (upper middle plot), radial glia-like and NPC-like areas. NPC enrichment is overlaid by the strong enrichment in the non-malignant areas. Predominantly, the hypoxic enrichment (right surface plot) and RG-like signature sharply separate the areas which were correlated to distinct metabolic patterns (right plot).

**Extended Data Figure 8:**
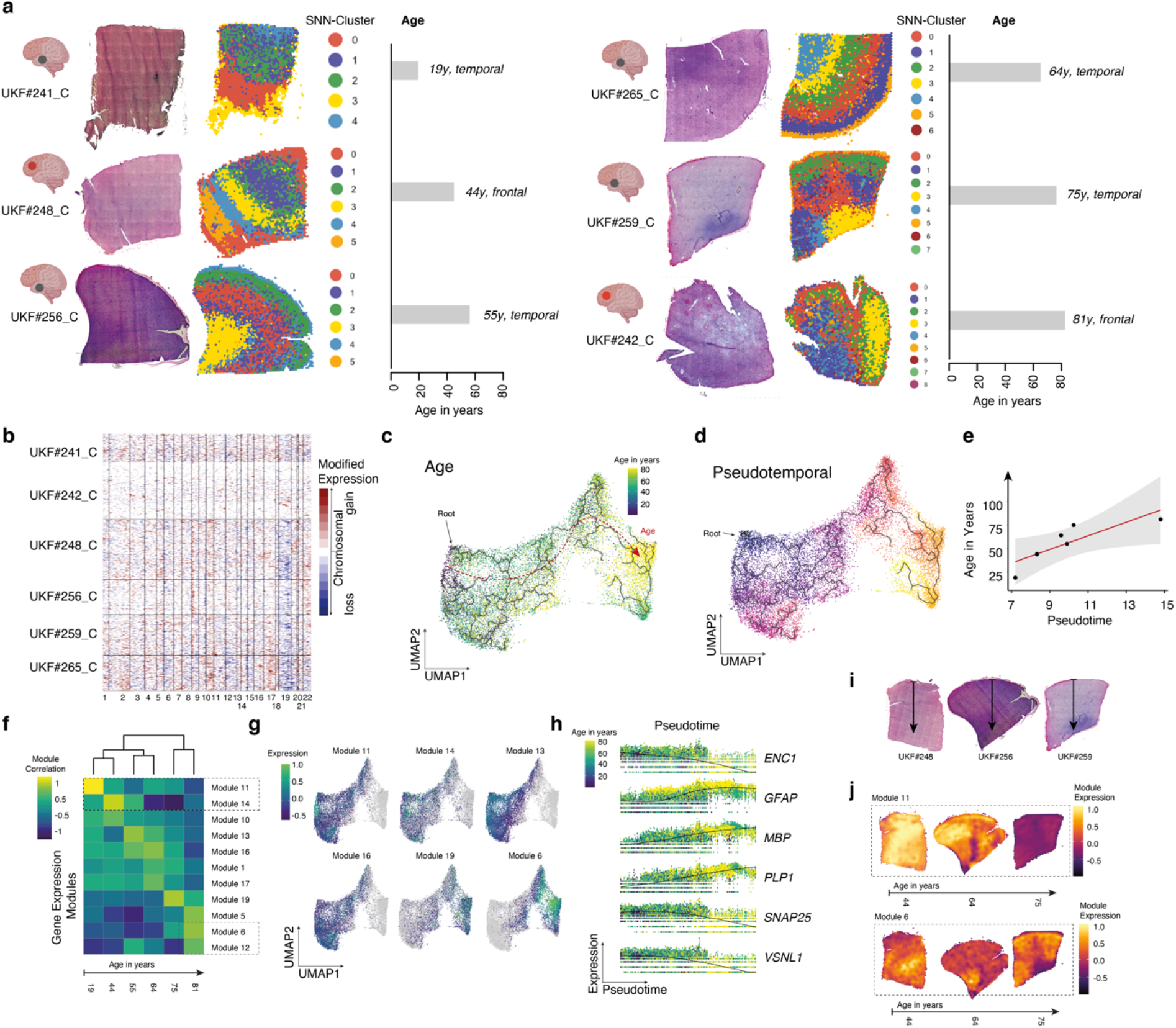
a) H&E staining (left upper) and surface plots with colored clusters (SNN-cluster approach) of non-malignant cortex samples. The correspondent age is given at the right side. b) CNV heatmap with gains in red and losses in blue indicate no CNV alteration in the collected samples. c-d) Dimensional reduction (UMAP) with colored age (left side, c) and pseudotime annotation (d). e) The pseudotime and real time (age patients) significantly correlate (R^2=0.67, p=0.031). f) Heatmap of age-related gene expression modules. g) Dimensional reducion (UMAP) with expression scores for age related modules. h) Gene expression of selected age-related genes. Spots are arranged along the pseudotime axis and colors indicate the age. I-j) Surface plots of gene expression scores of different modules.

**Extended Data Figure 9:**
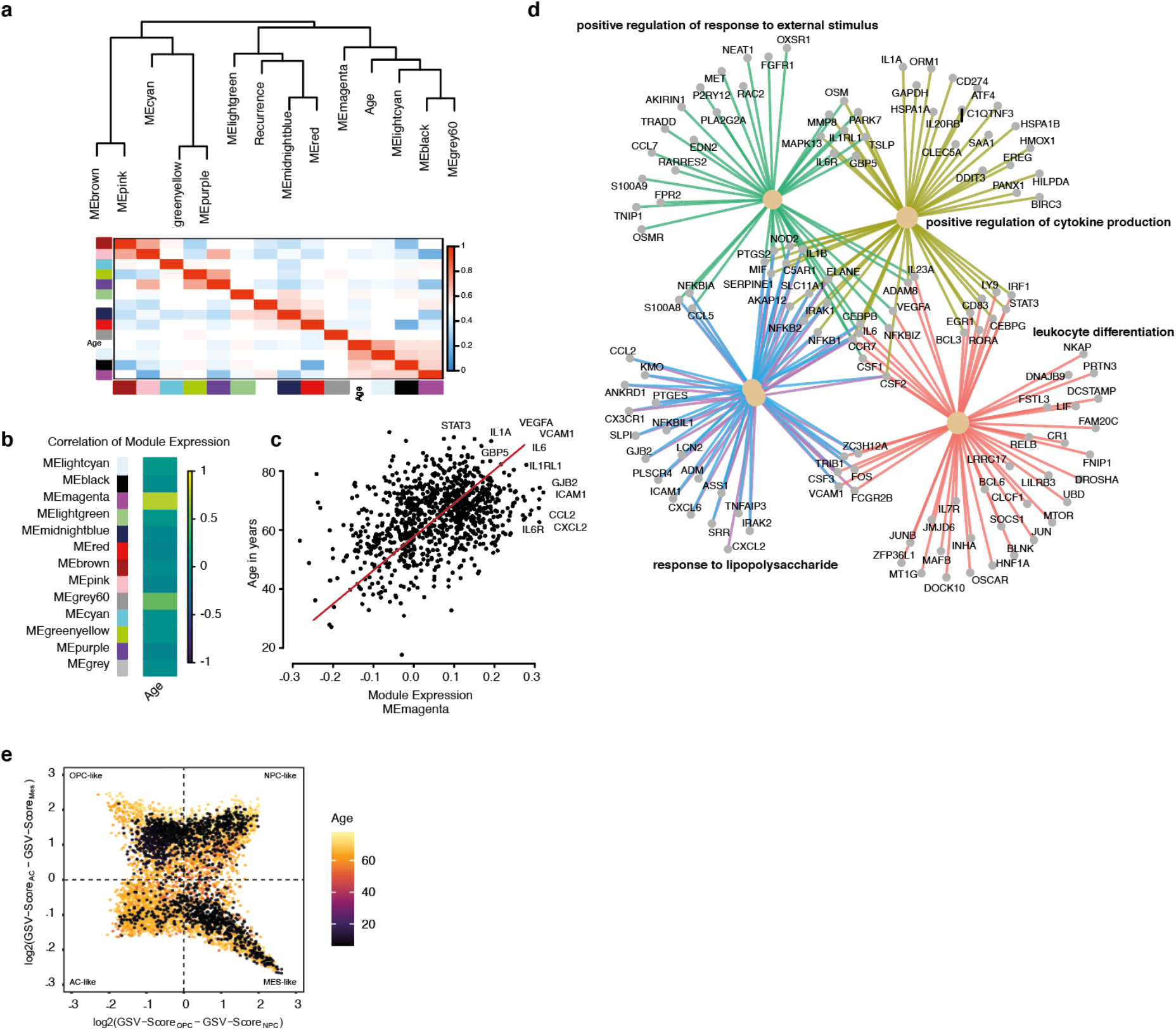
a-b) Weighted correlation network analysis of transcriptional data of the TCGA database. The analysis was designed to identify age-related gene expression signatures. Two modules were found to be significant associated with age (magenta and grey60) (b). c) Scatterplot of age (y-axis) and module expression (x-axis) with significant correlation (R^2 0.76 p<0.001). Top associated genes as printed. d) GSEA of genes (module magenta) confirmed a strong correlation of age and inflammatory gene expression signatures. e) Four-state scatterplot (Neftel et al.) indicate the four Neftel states based on the signature expression. The age of all patient in the Neftel dataset is annotated and colored.

**Extended Data Figure 10:**
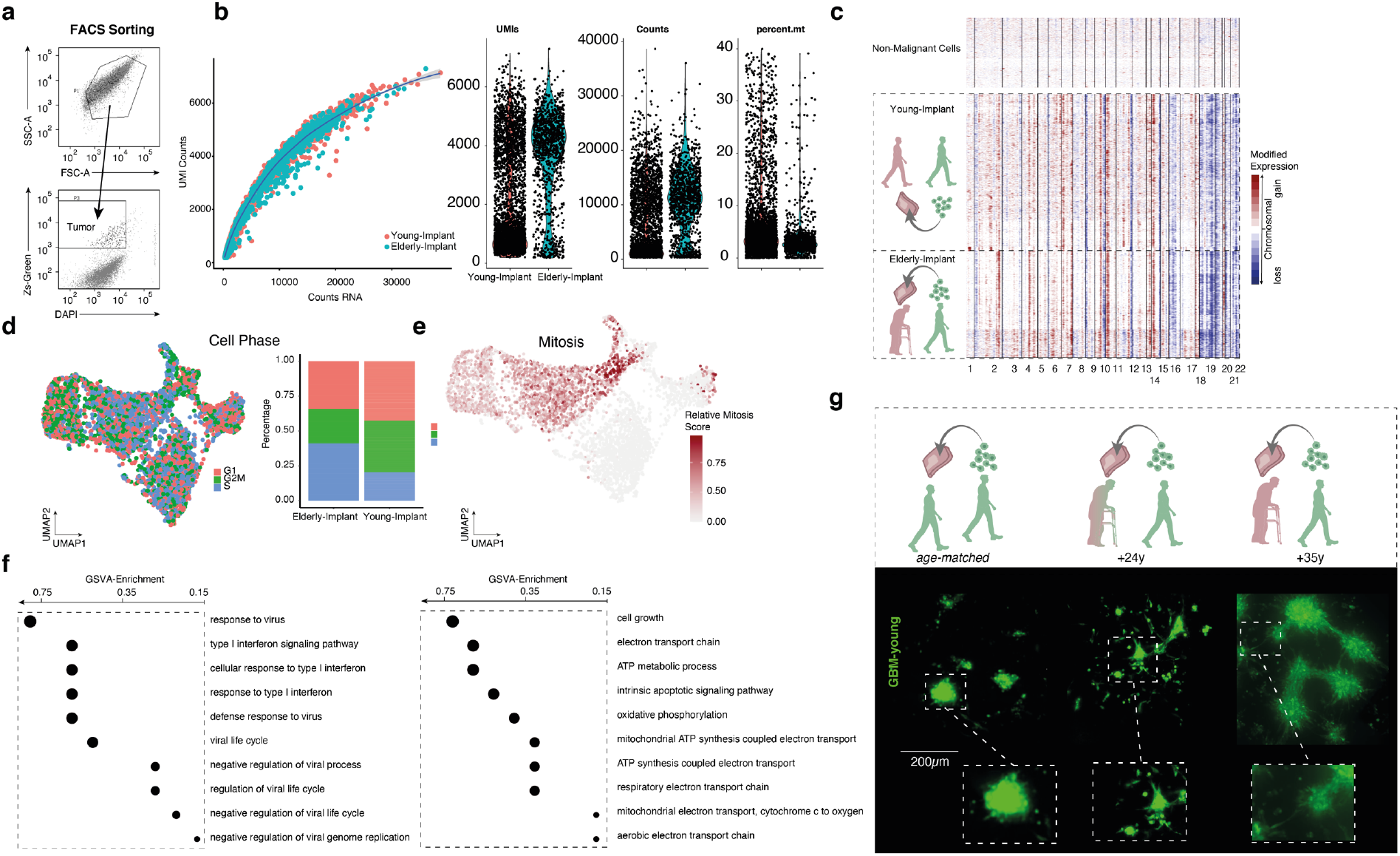
a) Scatter plots of used gate-strategy for cell sorting. b) Quality plots of the acquired scRNA-seq dataset. c) CNV plot of all cells, sharply separating between tumor and non-malignant cells. d) Dimensional reduction (UMAP) of separated tumor cells (cell phase plot) and correspondent fraction of cell phases between both sample sets. e) Dimensional reduction (UMAP) with colored cycling cells (Mitosis score). f) Enrichment analysis of genes highly differently expressed between both sample sets. g) Staining’s of slices with injection of “young tumor cells” (38 years) in slices (n=3) from different age groups. Tumor formation was highly different with a maximum growth in elderly cortex samples.

## Supplementary Table

Supplementary Table 1: Tissue Type (Macroscopic): T: Tumor, C: Cortex, TC: Tumor Core: TI: Tumor Infiltrative region

Supplementary Table 2: Gene sets from gene expression modules

Supplementary Table 3: Gene sets from pattern analysis

